# Trem2 Promotes Foamy Macrophage Lipid Uptake and Survival in Atherosclerosis

**DOI:** 10.1101/2022.11.28.518255

**Authors:** Michael T. Patterson, Maria Firulyova, Yingzheng Xu, Courtney Bishop, Alisha Zhu, Patricia R. Schrank, Christine E. Ronayne, Gavin Fredrickson, Ainsley E. Kennedy, Nisha Acharya, Xavier Revelo, Ingunn Stromnes, Tyler D. Bold, Konstantin Zaitsev, Jesse W. Williams

**Affiliations:** Center for Immunology, University of Minnesota, Minneapolis, MN USA; Department of Integrative Biology and Physiology, University of Minnesota, Minneapolis, MN USA; ITMO University, Saint Petersburg, Russia; Almazov National Medical Research Centre, Saint Petersburg, Russia; Department of Medicine, University of Minnesota, Minneapolis, MN USA; Department of Microbiology and Immunology, University of Minnesota, Minneapolis, MN USA

## Abstract

Atherosclerotic plaque formation is driven by the continued expansion of cholesterol loaded ‘foamy’ macrophages within the arterial intima. Foamy macrophages are primarily derived from newly recruited monocytes, but factors regulating monocyte specification toward foamy macrophage differentiation and prolonged survival in plaque remain poorly understood. We used trajectory analysis of integrated single cell RNA-seq data, along with a genome-wide CRISPR screening approach to identify Triggering Receptor Expressed on Myeloid Cells 2 (Trem2) as a candidate regulator for foamy macrophage specification. Loss of Trem2 led to a reduced ability of foamy macrophages to take up additional oxidized low density lipoprotein (LDL) cholesterol in vitro. Competitive chimera experiments showed that Trem2-deficient macrophages were less competent to form foamy macrophages when competed against Trem2-sufficient macrophages in vivo. In addition, myeloid specific conditional deletion of Trem2 resulted in a dramatic attenuation of plaque progression, even when targeted in established atherosclerotic lesions. This was independent of changes in circulating inflammatory cytokines, monocyte recruitment, or serum cholesterol levels, but due to a reduction in plaque macrophage proliferation and enhanced cell death. Mechanistically, we link Trem2-deficient macrophages with an inability for cells to sense cholesterol loading and failure to upregulate efflux molecules. Accumulation of cholesterol in the endoplasmic reticulum enhanced activation of the ER-stress response that increased susceptibility for cholesterol-toxicity and cell death in foamy Trem2-deficient macrophages. Overall, this study identifies Trem2 as a regulator of foamy macrophage differentiation, atherosclerotic plaque growth, and as a putative therapeutic target for future intervention studies.

## Introduction

Despite improvements in cardiovascular disease (CVD) outcomes in the last few decades, CVD remains a leading cause of death^1^. Atherosclerosis is a chronic disease of the artery wall that forms plaque, which are one of the primary causes of CVD and is driven by hyperlipidemia and vascular inflammation. Atherosclerosis progression is mediated by deposition of pathogenic low density lipoprotein (LDL) cholesterol particles into the arterial intima that accumulates within plaque associated macrophages, termed foam (or foamy) cells^2^. Foamy macrophages store esterified cholesterol in small lipid droplets, giving cells a bloated morphology^3^. These cells constitute a major portion of the total cellular presence in the early atherosclerotic plaque in humans and mouse models. Importantly, accumulation of foamy macrophages within lesions is associated with increased necrotic core formation and risk for plaque rupture, which contributes to two of the leading causes of CVD-related deaths, myocardial infarction and stroke^4^. In early lesions, foamy macrophages derive from a resident pool of aorta intima resident macrophages (Mac^AIR^), but are replaced by recruitment monocytes as plaque progresses^5^. Importantly, work suggests that foamy cells derived from either tissue resident or peripheral monocyte origins share a similar transcriptomic gene signature^5^. In larger atherosclerotic lesions the contribution of monocyte recruitment gives way to local macrophage proliferation as a primary mechanism for foamy macrophage maintenance and expansion^6^. However, the mechanisms involved in the specification and maintenance of foamy macrophages in early or late atherosclerotic lesions have yet to be fully understood^4,7,8^.

Recent single cell RNA-seq (scRNA-seq) transcriptional analysis has identified previously unrecognized heterogeneity for macrophage populations within the atherosclerotic aorta^9^. One notable finding was the identification of genes associated with foamy macrophages compared to nonfoamy macrophages isolated from atherosclerotic plaque, which included Triggering Receptor Expressed on Myeloid Cells 2 (Trem2)^10–12^. Trem2-expressing foamy macrophages were transcriptionally distinct from Il1b-expressing inflammatory macrophages, suggesting that divergent pathways may exist to regulate their differentiation or survival within atherosclerotic lesions^10^. Recent work has also found a correlation for the level of soluble Trem2 (sTrem2), which is cleaved from the macrophage cell surface, in the serum with cardiovascular-related mortality in high-risk patients^13^. Together, these studies have established Trem2 as a gene associated with foamy macrophage identity and lesion instability, but to date, no studies have addressed the function for Trem2 in atherosclerosis.

Trem2 is a cell surface lipid-sensor that plays a regulatory role in microglia function^14–16^, and polymorphisms in the Trem2 gene are causative for early onset Alzheimer’s-like dementia^17,18^. Trem2 signals through the adaptor molecules Dap10/Dap12 to activate Syk, PI3K, AKT, and mTOR pathways which leads to a pro-survival and anti-inflammatory response^19–22^. As a result, Trem2 broadly regulates phagocytosis, autophagy, cytoskeletal remodeling, and metabolic programming of macrophages^23,24^. Recent reports suggest that Trem2 may play dual roles in microglia during different stages of Alzheimer’s disease pathogenesis^25^. In early disease, Trem2 plays a role in the recognition and clearance of Aβ ‘seeds’ prior to plaque formation in the brain. However, in late-stage Alzheimer’s disease Trem2 is suggested to mediate lipid clearance from microglia^26^. In the periphery, Trem2 has been proposed to regulate homeostatic functions of adipose resident macrophages, and loss of Trem2 resulted in enhanced inflammatory signature and adipose hypertrophy^27^. Similarly, Trem2 expression on liver macrophages has been associated with a reparative phenotype and sTrem2, was proposed as a biomarker for liver disease pathogenesis^28^. Importantly, Trem2^-/-^ animals have systemic lipid dysregulation and elevated stress hormones, making the total body knockout mouse difficult to interpret for atherosclerosis or chronic inflammation models^27,29,30^. Overall, published studies support a potential role for Trem2 as a lipid-sensor and strong candidate for regulation of foamy macrophage function.

In this study, we performed trajectory analysis on integrated scRNA-seq data derived from murine atherosclerotic samples to define genes associated with monocyte differentiation toward foamy or nonfoamy plaque macrophage lineages. These data were compared against a genome-wide CRISPR inhibition screen that identified Trem2 as a regulator of foamy macrophage oxLDL uptake. Conditional deletion of Trem2 in a mouse model of atherosclerosis led to enhanced foamy macrophage death, reduced plaque macrophage proliferation, and reduced plaque lesion size. Trem2-deficient foamy macrophages showed an inability to down-regulate cholesterol biosynthesis pathways following lipid loading and reduced cell proliferation pathways. This was further associated with an upregulated endoplasmic reticulum (ER) stress response and enhanced macrophage cytotoxicity following cholesterol loading. Overall, this study reveals a new regulatory module in foamy macrophages that relies on Trem2 for regulating cholesterol accumulation and cell survival, and identifies Trem2 as a potential therapeutic candidate for future atherosclerosis therapy.

## Materials and Methods

### Animals

Mouse strains used for this study include; B6 (C57BL/6, Jackson Laboratory (Jax) 000664), Ldlr^-/-^ (B6.129 S7-Ldlr^tm1Her^/J, Jax 002207), CX3CR1^creER^ (B6.129P2(C)-Cx3cr1^tm2.1(cre/ERT2)Jung^/J, Jax 020940), Trem2^-/-^ (developed and provided by Dr. Colonna, Washington University)^31^, Trem2^flox^ (B6(C3)-Trem2^tm1c(EUCOMM)Wtsi^/AdiujJ, developed and provided by Dr. Lamb at Indiana University, Jax 029853)^17^, R26^tdTomato^ (B6.Cg-Gt(ROSA)26Sor^tm9(CAG-tdTomato)Hze^/J, Jax 007909), LysM^cre^ (Lyz2, B6.129P2-Lyz2^tm1(cre)Ifo^/J, Jax 004781). All mice are on the C57BL/6 background and bred in specific pathogen-free animal facilities maintained by the University of Minnesota (UMN) Research Animal Resources (RAR). When possible, littermates were used for experiments. Facilities were maintained at ~23°C with a 12/12 hour light/dark cycle. Cages were changed weekly and water was freely available through Lixit valve. All experiments and procedures were approved by the UMN Institutional Animal Care and Use Committee (IACUC). Experiments were performed in male and female mice in equal numbers. Experiments did not reveal significant differences between sex responses, therefore all datasets include combined data.

### HFD/TAM Diet feeding

In all experiments, animals were fed ad libitum. HFD (diet no. TD.88137; adjusted calories diet, 42% from fat) and TAM/HFD (diet no. TD.130903; adjusted calories diet, 42% fat, tamoxifen-citrate 400 mg/Kg) were purchased from Envigo Teklad. Mice were enrolled in studies between age 7-8 weeks and continuously maintained on HFD through the course of experiments, typically 8 or 16 weeks as described. In conditional Trem2 deletion experiments, aged-matched littermate CX3CR1^creER/+^ Trem2^fl/fl^ Ldlr^-/-^ mice were used for the experimental group (Trem2^ΔMac^), whereas CX3CR1^creER/+^ Trem2^fl/+^ Ldlr^-/-^ or CX3CR1^+/+^ Trem2^f/f^ Ldlr^-/-^ mice were combined for the Control (Cntl) group.

### scRNA-seq Data Integration and Analysis

Raw files were downloaded from NCBI Sequencing Read Archive. The kallisto bustools (v.0.46.1) workflow was used for the quantification of each sample in each dataset^32^. The count matrices obtained from the kallisto bustools pipeline were used as input. For the preparation of the atherosclerotic meta dataset and samples integration, the Seurat package (v.3.1.5) was used^33^. Samples from each study were processed and integrated into study-related objects which are available in Single Cell Navigator (https://artyomovlab.wustl.edu/scn/).

**Table 1:** scRNA-seq atherosclerosis studies from GEO database used in our study

| **Dataset ID** | **# samples** | **Mouse genetic background** | **Cell types** | **scNavigator link** |
| --- | --- | --- | --- | --- |
| GSE116240 (K. Kim et al. 2018) | 2 | *Ldlr*<sup>-/-</sup>, *Apoe*<sup>-/-</sup> | Cd45+, foam cells | GSE116240 |
| GSE123587 (Lin et al. 2019) | 2 | *Ldlr*<sup>-/-</sup> | CX3CR1+ monocyte lineage | GSE123587 |
| GSE131776 (Wirka et al. 2019) | 18 | *Apoe*<sup>-/-</sup> | SMC, non-SMC | GSE131776 |
| GSE131777 (Wirka et al. 2019) | 4 | *Apoe*<sup>-/-</sup> | SMC, non-SMC | GSE131777 |
| GSE149069 (Wolf et al. 2020) | 2 | *Apoe*<sup>-/-</sup> | Cd45+ | GSE149069 |
| GSE154692 (Jeon et al. 2020) | 4 | *Apoe*<sup>-/-</sup> | Cd45+ | GSE154692 |
| GSE154817 (Williams et al. 2020) | 2 | *Ldlr*<sup>-/-</sup>, WT | Cd45+ | GSE154817 |
| GSE155513 (Pan et al. 2020) | 14 | *Ldlr*<sup>-/-</sup>, *Apoe*<sup>-/-</sup> | SMC, non-SMC | GSE155513 |

#### Seurat analysis

Droplets with ambient RNA and noisy cells were filtered using *EmptyDrops* function from DropletUtils R package, and then genes which expressed in less than 200 cells were removed^34^. The fraction of mitochondrial genes was calculated for every cell, and cells with a mitochondrial fraction which were more than the sample-specific threshold defined by the confidence interval were filtered out. All samples were normalized using *SCTransform* function^35^. We next processed the data and features were detected using *SelectIntegrationFeatures*^33^. A list with all samples as its elements was prepared for integration using *PrepSCTIntegration* and *FindIntegrationAnchors* functions. Finally, samples were integrated using the *IntegrateData* function. Principal component analysis was run on the integrated object. For two-dimensional visualization of object structure, both tSNE and UMAP approaches were implemented using the first 20 principal components. For clustering purposes, functions *FindNeighbors* and *FindClusters* were used.

#### Identification of plaque monocytes and myeloid macrophages

All clusters were manually annotated using canonical gene markers. T cells (*Cd3d+*), B cells (*Cd79a+*), smooth muscle cells (*Sparc+*), proliferating cells (*Mki67+*), monocytes (*Treml4+, Ly6c2+, Sell+*) and different macrophages subtypes (*Retnla+, Adgre1+, Lyve1+, Fabp5+*) were identified in the prepared meta dataset. We first identified and removed the barcodes from T cells, B Cells, smooth muscle cells as well as proliferating cells. Then we used the expression of markers (*Lyve1+*, and *Mrc1^high^*) to separate adventitia macrophages from intima macrophages. The remaining barcodes (assumed to be monocytes and intima macrophages) were later reanalyzed from the very beginning (using the same steps as outlined above). Populations were found to be monocytes, intima macrophages, and dendritic cells. Cells that correspond to monocyte / macrophage populations were extracted and fully reanalyzed using Seurat (using the same steps outline above).

#### Trajectory analysis

All cells from the object which contained monocytes and intima macrophages were used, and *infer_trajectory* function from the dyno package (version 0.1.2) was used on the normalized counts (integrated assay, data slot) with the available slingshot singularity container (version 1.0.3)^36,37^. Trajectory visualization was implemented after dimensionality reduction by UMAP using the *dimred_umap* function. We also used dyno package to calculate gene importance scores for foamy or inflammatory differentiation along pseudotime^36^. In brief, the scores are calculated using a random forest regression model trained on gene expression values to predict pseudotime values. This allows easy interpretation, one can look at which genes are more important for the accuracy of such a model.

#### Differential expression across pseudotime

To find genes that are differentially expressed across pseudotime, the TradeSeq package (v.1.4.0) was used^38^. Raw counts (RNA assay, counts slot) were used as input expression and the design matrix corresponding to the study was used as fixed effects in order to remove the batch effect. We used *earlyDETest* to identify genes that are differentially expressed early after the branch point.

### Bone Marrow Chimera

Ldlr^-/-^ mice (recipients) were lethally irradiated with 1,100 rad using an X-ray irradiator using a split dose (550 rad each) provided 5 hours apart. Mice were rested for 4 hours, then adoptively transferred with donor bone marrow. Donor bone marrow cells (5×10^6^) were injected intravenously (i.v.) in a 100 μL volume to irradiated recipient mice. Mice were allowed to reconstitute for 8 weeks, then transitioned to HFD for atherosclerosis studies.

### Serum cytokine and cholesterol analysis

Blood was rested to clot at room temperature for 1 hour, then samples were centrifuged at 2,700 RPM for 10 minutes at 4°C in a tabletop centrifuge (Beckman Coulter). The supernatant (serum) was collected and assessed for cytokines and cholesterol content. Total cholesterol analysis was performed using the Wako/Fujifilm Cholesterol-E kit (#999-02601), following manufacturer’s instructions. Multiplex cytokine analysis was performed using LEGENDplex murine inflammatory panel (Biolegend #740446), following manufacturer’s protocol and analysis pipeline.

### In vitro culture

BV2 cells were a kind gift from Dr. Herbert Virgin (Washington University in St. Louis). BV2 cells were cultured in non-treated 10 cm tissue culture plates (VWR, #10062-880) in 5% fetal bovine serum (FBS, Corning, source from United States with low endotoxin) in Dulbecco’s Modified Eagle’s Medium (DMEM, Sigma-Aldrich, #D0819), with addition of 1% Penicillin-Streptomycin (Sigma-Aldrich, #P4333), HEPES (Sigma, #H0887), and MEM non-essential amino acids (Sigma-Aldrich, #M7145). Peritoneal cells were collected by lavage with Hanks Balanced Salt Solution (HBSS) with 2% FBS and 2mM EDTA as previously described^39^ and grown overnight in the same media as BV2 cells for assays. For foamy cell formation assay, cells were plated at 1×10^6^ cells in a 24-well plate (VWR, 10861-558) and treated overnight with 20 μg/mL of soluble cholesterol. Soluble cholesterol was generated as previously described^40^. In brief, methyl-beta-cyclodextrin was incubated with cholesterol at a 1:6 ratio, then stored at −20°C. DiI-labeled oxLDL (Kalen Biomedical, medium oxidized, #770282-9) was added to BV2 or peritoneal macrophages at 10 μg/mL for 4 hours. Cells were recovered from plates with a 3 minute incubation with 0.25% trypsin EDTA (Sigma-Aldrich, T4045), then scraped with rubber policeman to lift cells, washed with media, and stained for analysis.

### Generation of cas9 BV2 cell line

BV2 cells were transduced by lentivirus with Cas9 (pLX_311cas9) and selected with blasticidin (2 μg/mL) for 10 days. A single clone was isolated by dilution cloning. Cas9 expression was validated by flow cytometry using intracellular staining of cas9 protein with anti-cas9 monoclonal antibody (7A9-3A3, Cell Signaling, cat #48796). Cas9 activity was assessed by transducing the cas9-BV2 cells with pXPR_047, which expresses eGFP and a small guide RNA (sgRNA) targeting eGFP. Following transduction, cells were selected with puromycin (2 μg/mL) for 8 days. The percentage of GFP positive cells was assessed by flow cytometry on LSR II/Fortessa.

### Library preparation

The mouse Gouda (CP0074^41^) genome-wide CRISPR knockout library containing two sgRNAs per gene (purchased from Broad Institute), was lentivirally transduced into 9×10^7^ cas9-BV2 cells at a low MOI to result in approximately 30% of the cells infected (with approximately 1 sgRNA per cell) and achieve 500-fold coverage after puromycin selection. At 24 hours posttransduction, infected cells were selected with puromycin (2 μg/mL) for 5 days.

### CRISPR Screen

BV2 cells containing the gouda library were treated with 20 μg/mL soluble cholesterol overnight to generate foamy macrophages. Cells were then pulsed for 4 hours with DiI-oxLDL (Kalen). Cells were then sorted for oxLDL uptake, as indicated by DiI-lableing. The high and low 9% of DiI labeled cells were selected and then lysed for guide sequencing. Genomic DNA (gDNA) was purified and guides were sequenced with directed primers at the Broad Institute. Data were analyzed using Model-based Analysis of Genome-wide CRISPR-Cas9 Knockout (MAGeCK) algorithm^42^, raw read counts were first median-normalized to harmonize sample variations in regards of library size and count distribution. Next, a negative binomial approach involving mean-variance modeling was applied to determine the sgRNA abundance difference between control and test group. Subsequently, statistical scores are calculated and used to rank sgRNAs using MAGeCK test. We chose positively ranked genes as they represent the test (DiI Low) group. Log-fold change values reported by MAGeCK are used to perform pathway analysis using the MAGeCK pathway function.

### Generation of Trem2^-/-^ BV2 cell line

Targeted deletion strategy was adapted from prior work^43^. Cas9-BV2 cells were transfected with sgRNA targeting Trem2 (sgRNA sequence: CGTGTGTGCTCACCACACGC). To prepare the transfection, sgRNA was placed into the RNA backbone (pRDA_118) by mixing 8 μg guide RNA backbone with 2 μL BsmBI enzyme, and 5 μL NE Buffer 3.1 in a total volume of 50 μL in water and incubated at 55°C overnight. After incubation, 1 μL (10,000U/mL) CIP was added and incubated for 1 hour at 37°C. Digested sample was run on a 1% agarose gel and selected for ~8 kilobase size. The band was cut from the gel and purified using gel extraction kit (Qiagen, 28704). DNA was annealed and phosphorylated using forward and reverse oligos (CACCGCGTGTGTGCTCACCACACGC, AAACGCGTGTGGTGAGCACACACGC) in a ramp PCR setup increasing temperature by 0.1°C per second. The sample was then ligated with sgRNA overnight using T4 ligase. Stbl3 *E. Coli* were transformed by adding 5 μL ligation reaction mix to 50 μL of *E. coli* cells. Cells were rested on ice for 30 minutes, then cells were heat shocked at 42°C for 30 seconds and returned to ice. Cells were transferred to growth media and left in a shaking incubator for 1 hour at 37°C. 100 μL of cells were spread on an LB agar plate with 200 μg/mL ampicillin for selection. gRNA clones were selected after 24 hours and expanded in culture. Plasmids were isolated (Qiagen Miniprep kit, 27106) and sequenced for appropriate insertion. Zymo PurePlasmid Miniprep kit (D4209) was used to isolate endotoxin-fee plasmids for clones that were carried forward in the assay.

Cas9-BV2 cells were plated at 3×10^5^ in a 6 well plate in DMEM (without P/S) + 2 μg/mL of blasticidin. Transfection was performed using warm TransIT-LTI (Mirus Bio, #MIR2304), then vortexed. 5 ug of plasmid DNA were placed in 250 μL OptiMEM media (Gibco, 31985062), then 7.5 μL of transit-LTI reagent was added to DNA/OptiMEM solution. Samples were incubated at room temperature for 20 minutes, then given to cas9-BV2 cells. Cells were incubated for 3 days. Puromycin was added to treatment wells and incubated for additional 5 days for selection in appropriate insertion. Deletion efficiency for Trem2 targeting sgRNA was validated by flow cytometry and TIDE analysis using Sanger sequencing of the pooled clones, verifying ~60% deletion efficiency. Individual Trem2^-/-^ clones were isolated by limiting dilution and expansion of single clone wells. Deletion of Trem2 was validated on individual clones by flow cytometry. Eight total Trem2^-/-^ clones were isolated, and two clones were selected for experimentation in this study.

### Bulk RNA sequencing collection and analysis

*BV2 cells were lysed for RNA isolated by* Trizol directly from plates following overnight treatment in media or cholesterol. Samples were submitted to the University of Minnesota Genomics Core for RNA isolation (Qiagen RNeasy kit) and sequencing using the NOVAseq platform. Raw data was processed using CHURP pipeline developed by the Minnesota Supercomputing Institute, which implemented and integrated Trimmomatic, HISAT2, SAMTools and featureCounts. Data was aligned to the Mus musculus GRCm38 (Ensembl release 102) mouse reference genome. Differential expression analysis was adopted from DEseq2 (v.1.32.0). Pathway analyses were performed using the fgsea package (v.1.18.0).

### Flow Cytometry

Single cell suspensions were filtered through 100 μm nylon mesh (McMaster Carr), then washed in FACS buffer (HBSS with 2% FBS and 2mM EDTA). Supernatant was discarded and cell pellets were stained for 30 minutes at 4°C, protected from light. Antibodies were stained at 1 mg/mL in a volume of 50 μL, unless a specified concentration was specified by the manufacturer. Conventional flow cytometry was performed on a BD LSRFortessa or a BD LSRFortessa X-20. Spectral cytometry were collected using a Cytek Aurora. All machines are supported and maintained by the UMN flow cytometry core facility. Data were processed in Flowjo (Tree Star) or Cytek Spectroflo software.

**Table 2.** Antibodies used in this study.

| **Target Antigen** | **Conjugate** | **Species/Isotype** | **Clone** | **Company** | **Catalog #** |
| --- | --- | --- | --- | --- | --- |
| Trem2 | APC | Rat IgG2B | 237920 | R&D Systems | FAB17291A |
| Trem2 | FITC | Rat IgG2B | 78.18 | eBioscience | MA5-28223 |
| CD68 | Purified | Rat IgG2a | FA-11 | Biolegend | 137001 |
| Ki67 | Purified | Rabbit IgG | SP6 | Abcam | ab16667 |
| CD45 | BV480 | Rat IgG2B | 30-F11 | BD | 566168 |
| CD11b | BV605 | Rat IgG2B | M1/70 | Biolegend | 101237 |
| Ly6G | BV785 | Rat IgG2B | 1A8 | Biolegend | 127645 |
| Ly6C | BV421 | Rat IgG2B | HK1.4 | Biolegend | 128031 |
| CD115 | PerCPCy5.5 | Rat IgG2B | AFS98 | Biolegend | 135525 |
| TCRβ | APC | Hamster IgG | H57-597 | Biolegend | 109211 |
| CD19 | FITC | Rat IgG2B | 1D3 | Biolegend | 152403 |
| sXBP1 | AF647 | Rabbit IgG | E9V3E | Cell Signaling | 38139S |

### Cytotoxicity Assay

Cell viability was assessed using an LDH cytotoxicity detection kit (Sigma Aldrich) per the manufacturer protocol. Briefly, WT or Trem2^-/-^ BV2 cells were plated in 96 well plates and cultured overnight either in media alone or media with 20 or 40 μg/mL of cholesterol. The next day, supernatants were transferred to a new 96 well plate, along with wells of lysed cells and media alone for positive and negative controls. Samples were then treated with 100 μL of the reaction mixture and incubated at room temperature for 20 minutes. Then, samples were treated with 50 μL of stop solution and absorbance was immediately recorded on a Spectramax plus plate reader at 492 nm. Percent cytotoxicity was determined using the absorbance values minus the background controls and normalized to baseline (%Cytotoxicity= (sample value-negative control)/ (positive control-negative control)*100).

### Heart imaging/Immunofluorescence

Hearts were either fixed overnight in 4% paraformaldehyde/30% sucrose solution in 1x PBS or directly embedded. Hearts were embedded in OCT and frozen, then sectioned on a cryostat at 10 μm thickness. For staining, slides were warmed to room temperature for 10 minutes on the bench top. Samples were briefly fixed in 4% paraformaldehyde for 2 minutes, then washed with 1x PBS. Samples were blocked with 5% donkey serum and permeabilized with 1% triton x-100 for 30 minutes. Samples were washed two times with 1x PBS. Primary antibodies were diluted 1:500 in PBS and stained the samples for 1 hour. Samples were washed three times with 1x PBS, then stained with secondary antibodies conjugated to fluorochromes (1:1,000 dilution) for 30 minutes. Samples were washed 3x, then mounted with flouromount (Southern Biotec) and imaged. Samples were imaged using a Leica SP8 inverted confocal microscope (fluorescence imaging) or with an attached bright field light source and camera. Plaque area was measured using image J analysis tool, where blinded researchers identified areas of interest manually.

### Aorta Plaque Analysis

Aorta were surgically removed from mice and fixed in 4% PFA overnight at 4°C. Periadipose tissue was removed manually under a dissecting microscope (Leica S9i). Samples were then cut open and pinned en face to wax dishes. Samples were washed three times with water, then incubated in propylene glycol for 5 minutes. Next, the aortae were submerged in oil red O (Sigma O1516) for three hours, protected by light. Afterwards the dishes were washed in 85% propylene glycol for 5 minutes followed by three washes with water. Images were taken using Leica S9i stereo microscope with 10 megapixel color camera. Images were merged using Adobe photoshop and analyzed using Image J software. Plaque area were quantified by lab staff that were blinded to the sample identify.

### Human Plaques

Human plaque samples were isolated from the cranial circle of Willis and donated for research as part of the UMN anatomy bequest program. These tissues were collected post-mortem and de-identified with limited patient history. The plaque sample presented was isolated from a 93 year old female with a known history of cardiovascular disease. Samples were fixed in 4% formalin, dissected to approximately 1 mm size, then stained for CD68 and Trem2, and imaged by confocal microscopy in whole-mount. Images were analyzed using Bitplane Imaris v9 software.

### Statistical Analysis

Graphs were generated and statistical analysis performed in Graphpad Prism software. In general, comparison between two experimental groups utilized a student’s T-test, whereas comparisons of more than 2 groups utilized a two-tailed ANOVA analysis. Specific analysis is included in the figure legend of each experiment. P-value <0.05 were considered statistically significant.

## Results

### Trem2 is associated with foamy macrophage differentiation

Recent efforts in scRNAseq analysis approaches have generated high dimensional analysis of immune cell profiles of atherosclerotic plaques from mouse and human^44,45^. These studies have resolved fine detail in heterogeneity of cell subsets that were previously unknown. However, differentiation kinetics or genetic regulators of fate specification between key subsets, such as between monocytes and inflammatory or foamy macrophages, remains unclear. To enable accurate *in silico* identification of the differentiation trajectory from monocytes to macrophages, we created a meta-dataset of immune cells associated with mouse atherosclerosis. For this, we used eight publicly available scRNA-seq studies focused on atherosclerosis research using mouse models downloaded from NCBI Gene Expression Omnibus (Figure 1A). Some studies (K. Kim et al. 2018; Lin et al. 2019; Wolf et al. 2020; Jeon et al. 2020; Williams et al. 2020) are focused on the immune populations present in the plaque while others (Wirka et al. 2019; Pan et al. 2020) are focused on smooth muscle cell (SMC) phenotypic modulation and switch during the progression of atherosclerosis^5,10,46–50^. We uniformly processed all these datasets using raw data from NCBI Sequence Read Archive and integrated all the datasets together. Total clusters identified and differential gene expression analysis supported prior Meta-analysis studies (Supplemental Figure 1)^9^. To remove nonmacrophage clusters and cell clusters that were not present within atherosclerotic lesions, we used prior sequencing profiles to identify and enriched for the clusters corresponding to monocytes and intima macrophages^10^. The resulting dataset contained 4 main clusters (Figure1 B-C), which could be described using the canonical gene markers: monocytes (*Hp+ Treml4+ Ly6c2+ Sell+*), foamy macrophages (*Fabp5+ Mmp12+ Gpnmb+ Itgax+*), inflammatory macrophages (*Tnf+ Nlrp3+ Mgl2+*), and MHC-II^high^ macrophages (*MHC-II+ Cd74+*). Inflammatory macrophages in our classification share the expression of many MHC-II genes with MHC-II^high^ macrophages, yet, MHC-II^high^ macrophages lose most of the inflammatory signature. Importantly, all four intima-associated clusters were represented in each individual scRNA-seq dataset, suggesting high reproducibility for cluster identities across labs and animal models (Figure 1D). Furthermore, our analysis confirmed the rare representation of bona fide monocytes, which supports the necessity of integrating all available studies for trajectory studies since no single dataset was sufficient to generate fine detail between these key clusters.

**Figure 1.**
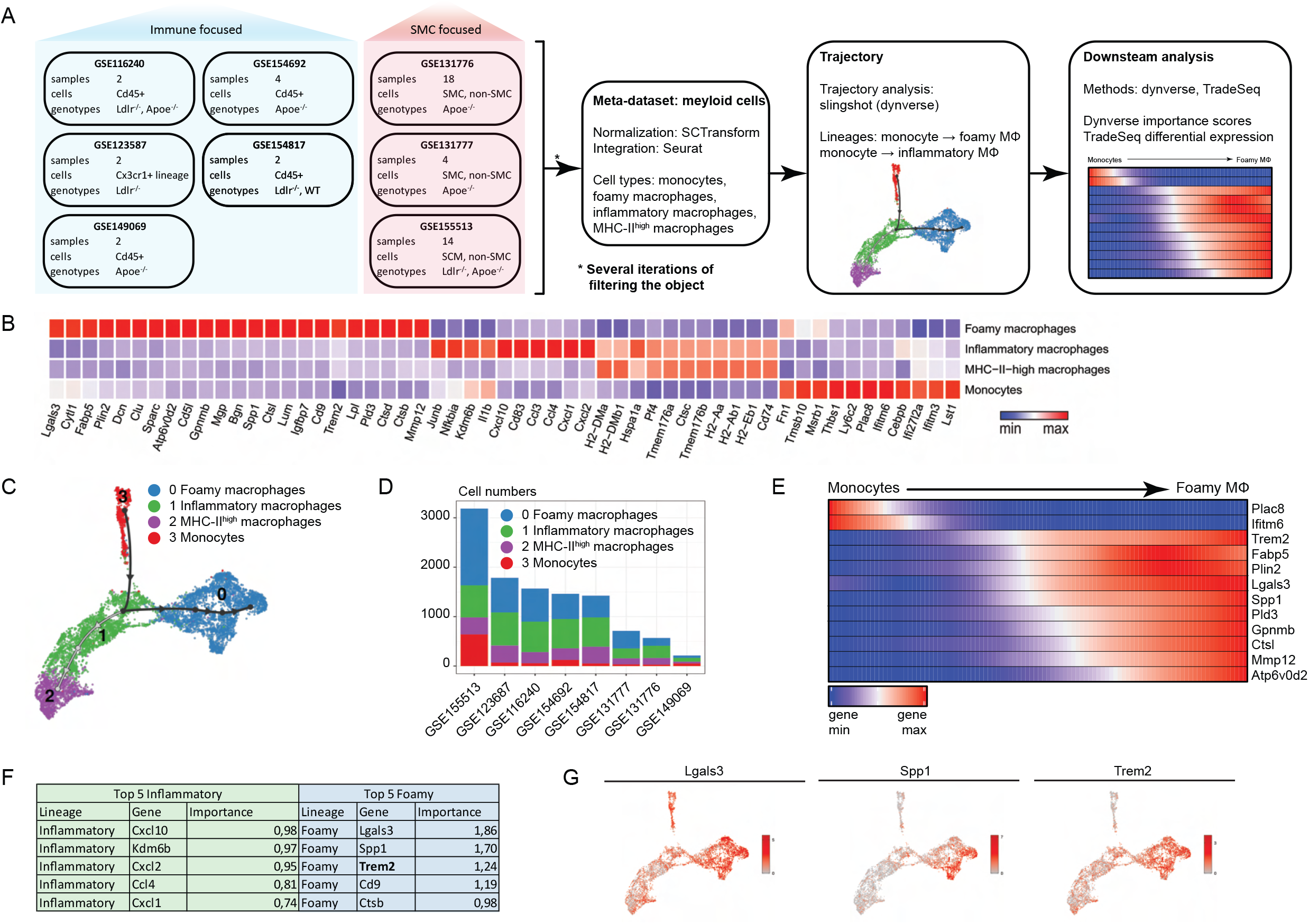
META scRNA-seq trajectory analysis identifies genes associated with foamy macrophage differentiation in atherosclerotic plaques. A) scRNA-seq datasets from atherosclerosis studies were integrated into a single metadataset. Based on cluster gene enrichment signatures cells were filtered to isolate intima-associated monocyte and macrophage clusters. Cells were examined for trajectory analysis and differential gene expression. B) Four main clusters of intima associated monocytes/macrophages were identified and annotated based on enriched gene signatures. Top differential expressed genes are displayed in association with the different clusters. C) Trajectory analysis was performed to determine the potential differentiation pathways used by foamy or inflammatory clusters. Data emphasize a monocyte origin and bifurcation toward terminal macrophage differentiation endpoints. D) Monocyte and macrophage cluster representation from original studies is displayed, emphasizing the presence of all clusters from each independent study. E) Pseudotime trajectory was plotted between monocyte (cluster 3, origin) and foamy macrophages (cluster 0, end point). Genes associated with monocyte lineage, including Plac8 and Ifitm6, were rapidly lost and genes associated with foamy macrophage specification were enriched across pseudotime. F) Trade-seq analysis algorithm predicted genes most likely associated with lineage commitment, called importance index. Top predicted genes for inflammatory and foamy differentiation are outlined in the table. G) The top three genes associated with foamy cell “importance index” were plotted on tSNE project map for gene expression profile.

To determine predicted differentiation trajectories between intima myeloid cell subsets, we applied the slingshot algorithm from the Dynverse^36,37^ to our integrated dataset. Using monocytes as the origin point, we reveal two trajectories that suggest binary fate determinates toward either foamy or inflammatory macrophage terminal lineages (Figure 1D). Surprisingly, trajectory analysis suggests that there is an intermediate inflammatory population that is shared between differentiation programs (Supplemental Figure 2A). This appears to be a shared program by newly recruited monocytes that upregulate modest levels of MHCII and other inflammatory associated genes prior to commitment toward foamy lineage, where inflammatory genes are downregulated, or inflammatory lineage, where inflammatory genes and chemokines are further upregulated from this intermediate transition stage (Supplemental Figure 2B). While it is difficult to meaningfully split the inflammatory macrophage cluster into several due to the shared continuous expression of many canonical inflammatory genes (Il1b, Tnf, Nlrp3), there were a series of genes that were reserved for macrophages that further committed to inflammatory fate, including MHC II genes, Ccl3, and Ccl4 (Supplemental Figure 2C-D).

Next, we sought to visualize the kinetics of the monocyte-to-foamy macrophage differentiation program on pseudotime ordering. Heat map shows the gene expression changes with pseudotime plotted on the x-axis for selected markers of foamy macrophages (Figure 1E). While the patterns of activation of foamy macrophage-associated genes *Lgals3, Spp1*, and *Trem2* are very similar, the heat map suggests earlier activation of *Trem2* transcript during lineage commitment. After we built the trajectories and obtained pseudotime ordering, we used Dynverse to also obtain gene importance scores associated with each terminal differentiation outcome^36^ (Figure 1F). Lastly, gene expression plots for top candidate genes, *Lgals3, Spp1*, and *Trem2* are shown to confirm specificity with the foamy macrophage cell cluster (Figure 1G). Absolute values of expression are also very different for these genes, *Lgals3* has a continuous expression from monocytes to foam cells, whereas expression of *Trem2* is only detected in intermediate inflammatory macrophages and is further upregulated in foamy macrophages. Overall, this analysis suggests that monocyte commitment toward fully differentiated plaque macrophages occurs as a binary fate-decision from a common inflammatory intermediate population. In addition, these data further identify candidate genes that may be key regulators for differentiation into these terminal states.

### Genome wide CRISPR-inhibition screen to determine regulators of oxLDL uptake

Single cell trajectory and differential gene expression analysis provided a detailed map of transcriptional changes that occurs during foamy macrophage differentiation. However, it is unable to define which genes regulate foamy macrophage differentiation. To determine whether genes expressed during foamy macrophage commitment could also influence the ability of macrophages to accumulate oxidized LDL (oxLDL), we designed an unbiased in vitro CRISPR screening approach. Using the BV2 myeloid cell line, we used lentivirus infection to insert Cas9 and the “Gouda” knockout pooled CRISPR-guide library containing 2x coverage across the genome^41^. We designed a CRISPR-screen to detect genes associated with oxLDL uptake by foamy macrophages. To do this, BV2-Gouda cells were loaded with soluble cholesterol overnight to generate lipid loaded macrophages, then pulsed with fluorescently labeled DiI-conjugated oxLDL particles (Figure 2A-B). After four hours, cells were collected and separated into DiI^low^ or DiI^high^ populations by fluorescence activated cell sorting (FACS). Approximately the top 9% and bottom 9% of cells labeling with DiI were sorted and lysed to sequence for CRISPR guide enrichment. Differential expression between DiI^low^ versus DiI^high^ cells revealed gene targets enriched for enhanced or reduced oxLDL uptake, including Trem2 (Figure 2C). Unbiased analysis of guide enrichment rank-ordered against p-values are shown (Supplemental Figure 3). Selected genes associated with lipid processing, classical activation, or alternative activation pathways are also shown (Figure 2D). Interestingly, of the top 15 “importance-index” genes identified in foamy cell trajectory analysis in Figure 1, Trem2 was the top enriched gene associated with the regulation of oxLDL uptake of foamy BV2 cells (Supplemental Figure 3B).

**Figure 2.**
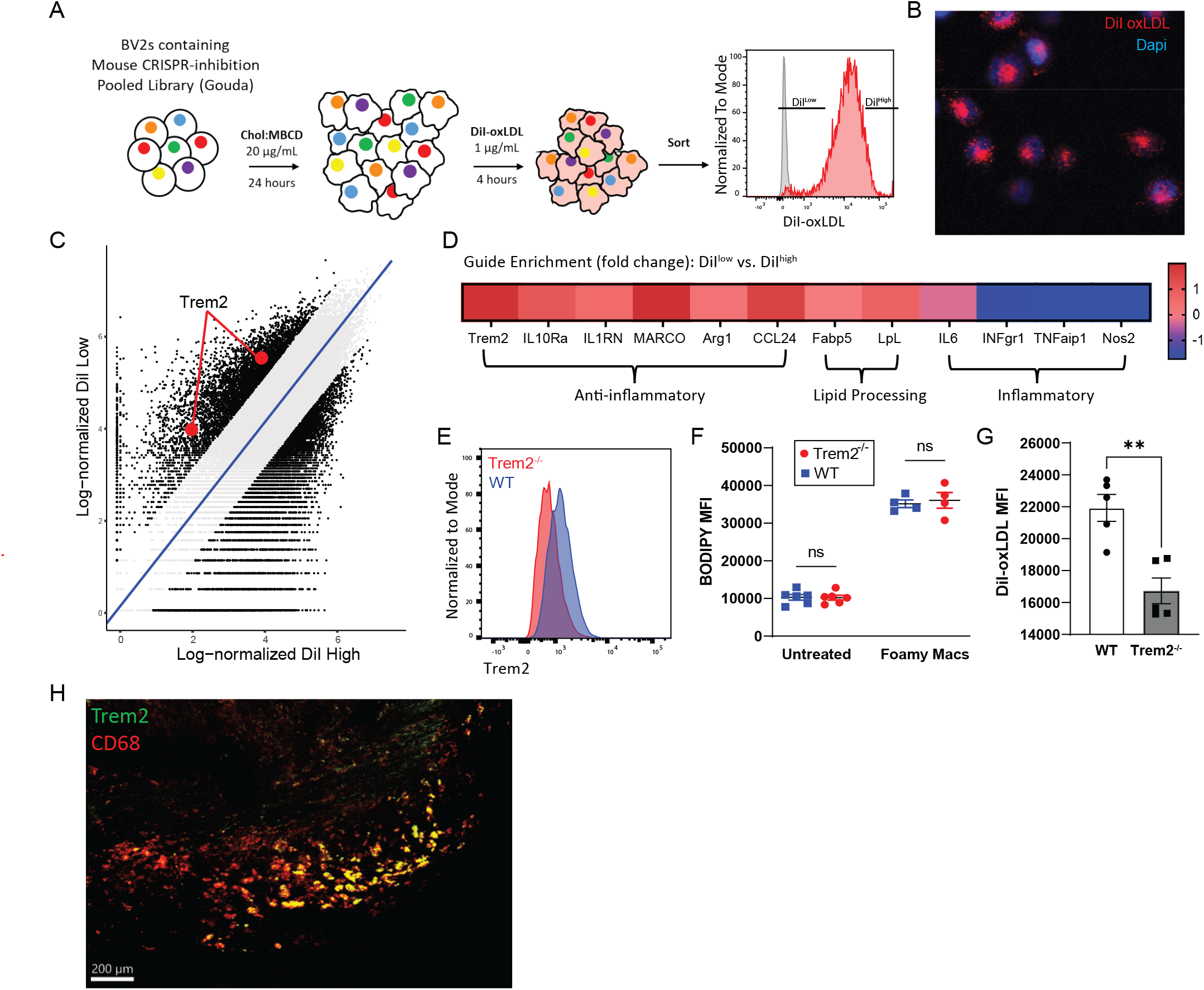
Genome-wide CRISPR-screen identifies Trem2 as a candidate regulator for foamy macrophage formation. A) Schematic for CRISPR inhibition screening approach for oxLDL uptake. BV2 macrophages were loaded with CRISPR pooled guide library (Gouda). Cells were made foamy by overnight treatment with soluble cholesterol, then challenged for 4 hours with DiI-oxLDL and sorted for DiI-high and DiI-low cells. Guides were sequenced from sorted populations. B) Confocal micrograph showing BV2 DiI uptake after 4 hour incubation with DiI-oxLDL. C) CRISPR-guide enrichment comparing log-normalized enrichment in DiI-high (X-axis) versus DiI-low (Y-axis). D) Selected gene enrichments comparing DiI-low vs. DiI-high. E) Peritoneal macrophages were isolated from WT or Trem2^-/-^ mice and treated with soluble cholesterol to induce foamy cell formation. After overnight culture, cells were analyzed for Trem2 expression by flow cytometry. F) Bodipy staining for total neutral lipid accumulation was performed by flow cytometry on peritoneal macrophages from WT or Trem2^-/-^ mice, cultured overnight in media alone or in media with soluble cholesterol. G) Peritoneal macrophages were isolated from WT or Trem2^-/-^ mice and treated with soluble cholesterol to induce foamy cell formation. After overnight culture, cells were treated with DiI-oxLDL for 4 hours and assessed for uptake by flow cytometry. H) Cranial artery plaques were stained for CD68 and Trem2 to identify co-expressing foamy macrophages within human plaques.

Next, we sought to validate the CRISPR screen result using ex vivo cultured primary macrophages. Peritoneal macrophages were isolated from C57Bl/6 (WT) or Trem2^-/-^ mice and treated with soluble cholesterol overnight. Overnight treatment of peritoneal macrophages was sufficient to induce Trem2 expression on WT peritoneal macrophages, as detected by flow cytometry (Figure 2E). Bodipy staining for total neutral lipids in cholesterol treated cells confirmed similar lipid accumulation in both WT and Trem2^-/-^ peritoneal-derived macrophages that were untreated or foamy differentiated (Figure 2F). However, DiI-oxLDL treatment resulted in reduced fluorescence in Trem2^-/-^ foamy macrophages compared to WT controls (Figure 2G), confirming a dependence of foamy macrophages on Trem2 for efficient oxLDL uptake. Overall, these data suggest that Trem2 regulates the ability of lipid loaded macrophages to take up additional oxLDL. Finally, Trem2 was previously found to be expressed at the transcript level by human plaque macrophages by scRNA-seq analysis^51^. We confirmed by protein immunofluorescence that Trem2 protein was expressed on foamy macrophages present in human atherosclerotic plaque (Figure 2H). Thus, Trem2 is shared by plaque-associated macrophages in mouse and human, and is a putative regulator of foamy macrophage formation.

### Trem2 is required for foamy cell formation in vitro and in vivo

Trem2 has been shown to regulate macrophage polarization, phagocytosis, and survival, but the role of Trem2 remains to be examined in models of atherosclerosis. Since Trem2^-/-^ mice are reported to have elevated cholesterol levels following high fat diet (HFD) feeding, we elected to utilize a mixed bone marrow chimera approach to normalize cholesterol levels between strains and allow for examination between WT and Trem2^-/-^ macrophages during atherosclerotic plaque progression. Atherosclerosis-susceptible Ldlr^-/-^ mice were lethally irradiated, then transplanted with a 50/50 mix of Trem2^-/-^ bone marrow cells or control bone marrow cells that express LysM^cre^ R26^tdTomato^ reporter allele to allow for identification of cell origin by fluorescence. Mice were rested for 8 weeks following irradiation to fully reconstitute the immune system, then chimeric mice were fed an HFD for 8 weeks to induce atherosclerotic plaque formation (Figure 3A). Blood analysis confirmed efficient mixing of Trem2^-/-^ (unlabeled) and control (tdTomato^+^) monocytes in the chimeric mice (Figure 3B). Confocal analysis of the aortic atherosclerotic plaques revealed a distinct enrichment of tdTomato^+^ cells in cells resembling foamy morphology (large, bloated) that co-stained for CD45 (white), whereas CD45^+^ tdTomato^-^ macrophages were largely associated with smaller and ramified morphology (Figure 3C). For quantification, foamy macrophages were blindly identified using morphology and CD68 antibody staining, then separated into tdTomato-positive (WT) or -negative (Trem2^-/-^) populations. Analysis revealed equal contributions to blood monocytes, but that Trem2^-/-^ macrophages failed to effectively compete against WT macrophages to differentiate into foamy macrophages in atherosclerotic plaque (Figure 3D). To confirm this finding, we also employed a “Foam FACS” approach to determine foamy macrophage formation by flow cytometry in our mixed bone marrow chimera mice^10^. After HFD feeding, fresh aorta were isolated, cleaned, and digested to liberate macrophages for flow cytometry. Cells were antibody labeled to identify macrophages (CD64^+^ CD11b^+^), then separated into Trem2^-/-^ (tdTomato^-^) or Control (tdTomato^+^) and cells were assessed for lipid content (bodipy) and side-scatter (ssc) as a measure of granularity (Figure 3E). Analysis showed that control macrophages were more effective at taking up lipid compared to Trem2^-/-^ macrophages (Figure 3F) and that a larger percentage of tdTomato^+^ cells were phenotypically foamy (Figure 3G). Together, these data support that Trem2 promotes the formation of foamy macrophages in atherosclerotic plaque.

**Figure 3.**
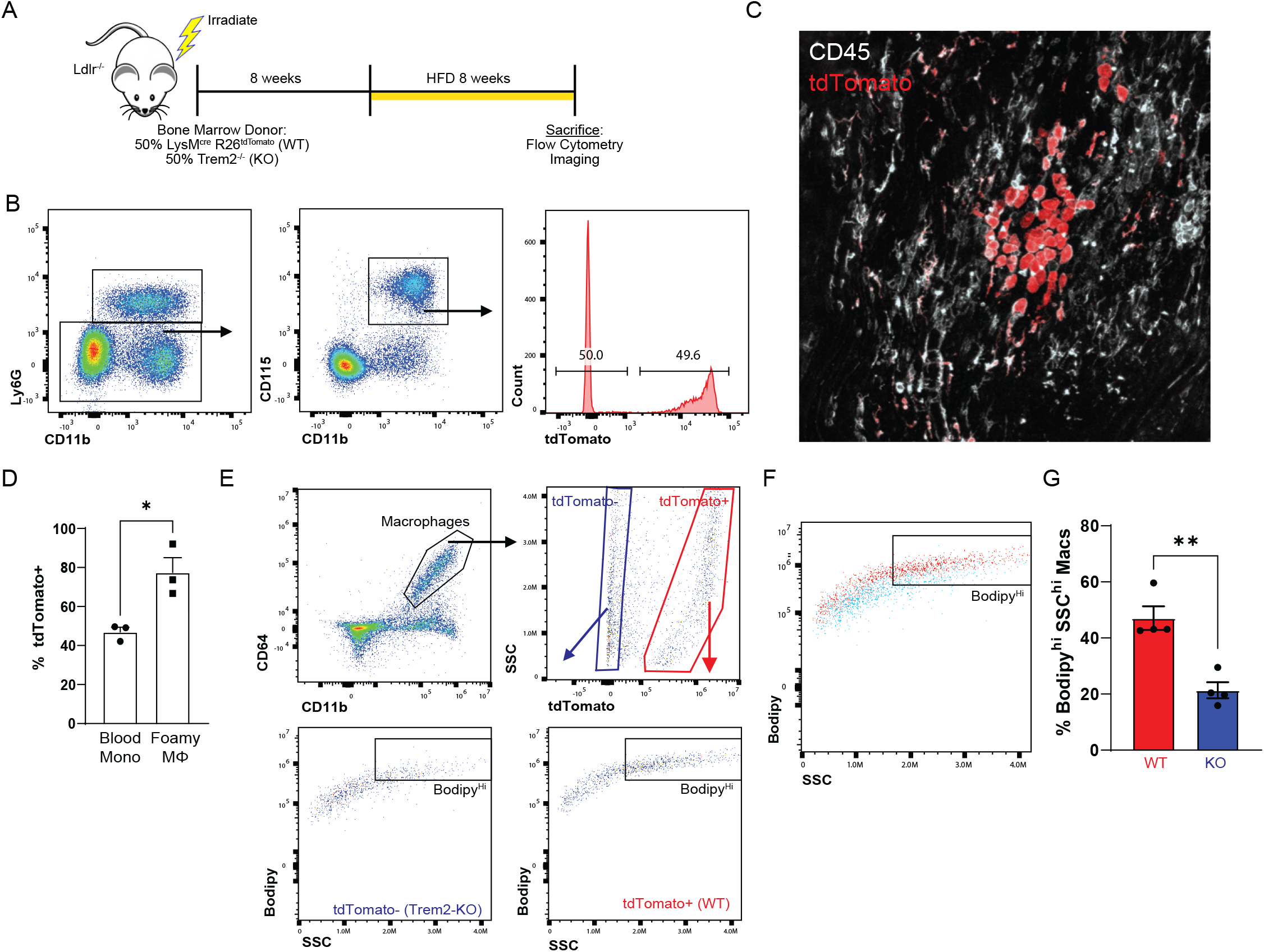
Trem2-deficient macrophages are outcompeted by WT macrophages to form foamy cells in atherosclerotic plaque. A) Schematic for mixed bone marrow chimera experiment. Ldlr^-/-^ mice were lethally irradiated and rescued by donor bone marrow from (50%) LysMcre R26tdTomato (WT) and (50%) Trem2^-/-^ mice. Recipient mice were rested for 8 weeks, then fed a HFD for additional 8 weeks to induce atherosclerosis. B) Flow cytometry gating of blood immune cells (CD45+) after 8 weeks HFD feeding, showing ratio of monocytes derived from WT (tdTomato+) and Trem2^-/-^ progenitors. C) Confocal micrograph of whole mount aorta showing tdTomato labeling (red) and CD45 (white) staining to define cellular contributions to foamy macrophages. D) Quantification of tdTomato+ cells in blood compared to foamy macrophages from whole mount aorta images. E) Foamy FACS was performed on CD64+ CD11b+ macrophages isolated from mixed bone marrow chimera aorta. Macrophages were separated into tdTomato+ and tdTomato-populations and then assessed for foamy representation by SSC and bodipy (neutral lipid) staining. F) Flow cytometric overlap between tdTomato+ (red) and Trem2^-/-^ (blue) derived macrophages from digested atherosclerotic aorta. G) Quantification derived from flow cytometric foamy FACS comparing relative contribution to foamy macrophages.

### Loss of Trem2 on macrophages attenuates atherosclerosis progression

Given the systemic defects associated with the total body Trem2^-/-^ mouse model, we crossed the Trem2 conditional knockout (Trem2^flox^) mouse with the CX3CR1^creER^ myeloid-specific inducible-Cre mouse on the Ldlr^-/-^ background (Trem2^ΔMФ^). This approach allows for deletion only on CX3CR1-expressing cells and temporal control to allow for normal development of tissues prior to the deletion of Trem2. Control (Cntl) animals for these experiments were littermates and included Cre-negative animals (CX3CR1^+/+^Trem2^fl/fl^ Ldlr^-/-^) and Cre-positive animals (CX3CR1^creER/+^ Trem2^fl/+^Ldlr^-/-^). Both control strains showed comparable results, thus were combined for analysis. To test the role of Trem2 in plaque formation, conditional knockout mice or controls were continuously fed a tamoxifen-enriched HFD (TAM-HFD) for 8 or 16 weeks to induce atherosclerosis and Trem2 deletion in CX3CR1-expressing cells. Strikingly, after 8 weeks of TAM-HFD feeding, atherosclerotic plaques in both the aortic arch and aortic sinus were dramatically reduced in the Trem2^ΔMФ^ mice compared to controls (Figure 4A-B). Importantly, this result was independent of changes in serum cholesterol or body weight (Figure 4C-D). Reduced atherosclerotic plaque formation in that aortic arch and aortic sinus of Trem2 conditional deletion mice were replicated at later atherosclerosis time points after 16 weeks TAM-HFD feeding (Figure 4E-F). Again, the reduction in plaque was not associated with changes in cholesterol or weight (Figure 4G-H). Together, these data show Trem2 is required in CX3CR1-expressing macrophage for atherosclerosis progression, and supports the hypothesis that Trem2 is a key regulator of foamy macrophage formation in atherosclerotic lesions.

**Figure 4.**
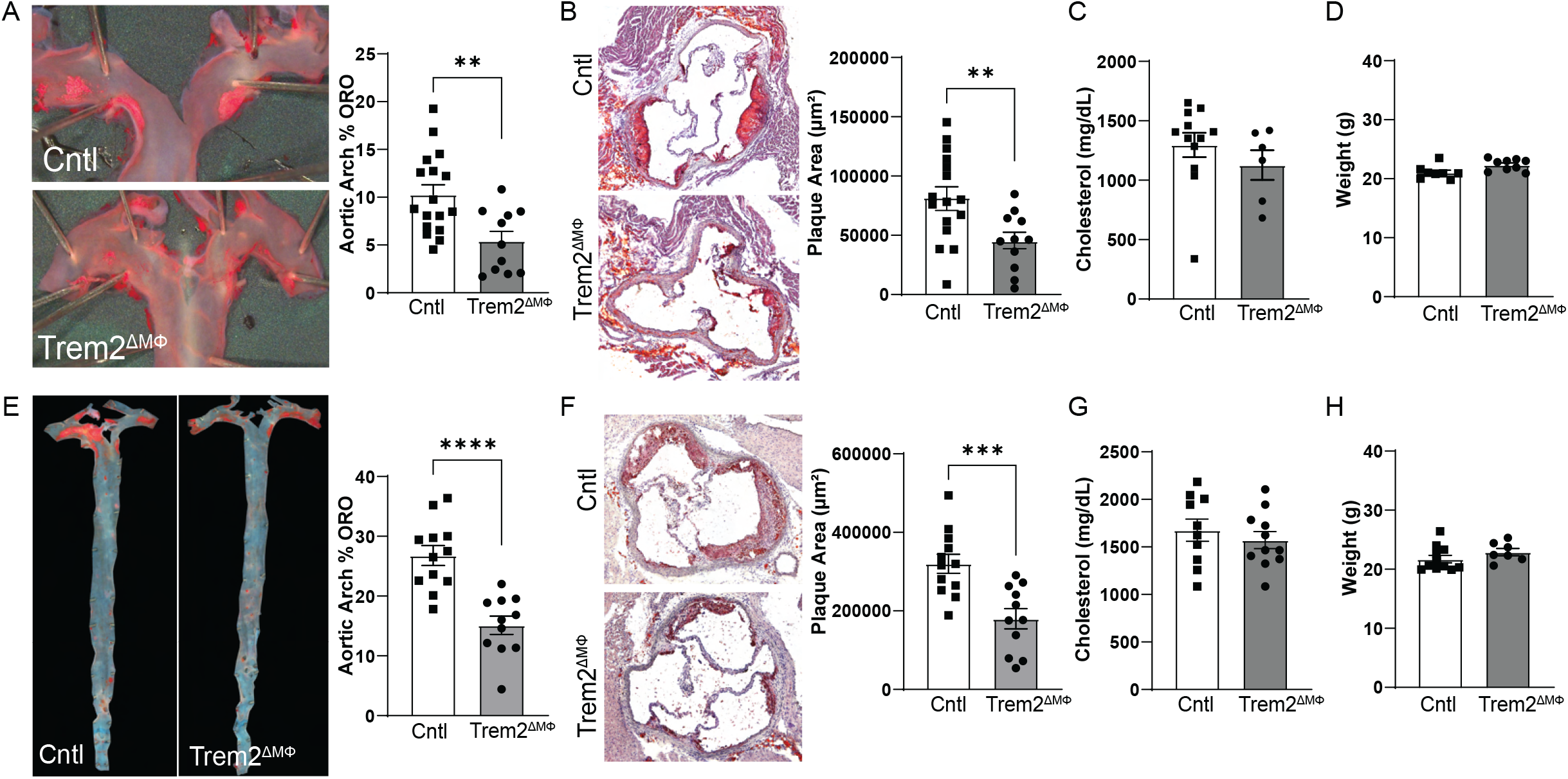
Conditional deletion of Trem2 on macrophages attenuates atherosclerotic plaque progression. CX3CR1^creER^ Trem2^flox/flox^ Ldlr^-/-^ or littermate control mice were fed a tamoxifen-enriched HFD (TAM-HFD) for 8-(A-C) or 16-weeks (D-F). A) After 8 weeks of TAM-HFD aorta were analyzed by en face analysis for percentage oil red O (ORO) staining on the arch. B) Aortic sinus plaque area measured after ORO staining in 8 week TAM-HFD samples. C) Serum cholesterol levels from 8 week TAM-HFD fed mice. D) Weight data from 8 week TAM-HFD fed mice. E) En face ORO staining of aorta following 16 weeks TAM-HFD feeding. F) Aortic sinus plaque area after 16 week TAM-HFD feeding. G) Serum Cholesterol after 16 week TAM-HFD feeding H) Weight of mice after 16 weeks TAM-HFD feeding.

### Trem2 regulates foamy macrophage survival in plaque

To determine mechanisms regulating plaque progression in Trem2^ΔMФ^ mice, we initially investigated whether there were systemic changes in inflammation. First, by serum cytokine multiplex assay, we observed no significant changes across a broad panel of 10 cytokines associated with atherosclerosis progression at 8 or 16 weeks (Figure 5A-B). Next, since increased blood monocyte numbers are associated with elevated atherosclerotic plaque formation^52^, we performed flow cytometry to assess whether there were systemic changes in peripheral blood immune cell populations (Supplemental Figure 4A). Data revealed no dramatic changes in monocyte or other immune cell numbers in the blood, suggesting that changes in systemic inflammation were not a major driver of the reduced atherosclerosis observed in Trem2^ΔMФ^ mice (Figure 5C, Supplemental Figure 4B).

**Figure 5.**
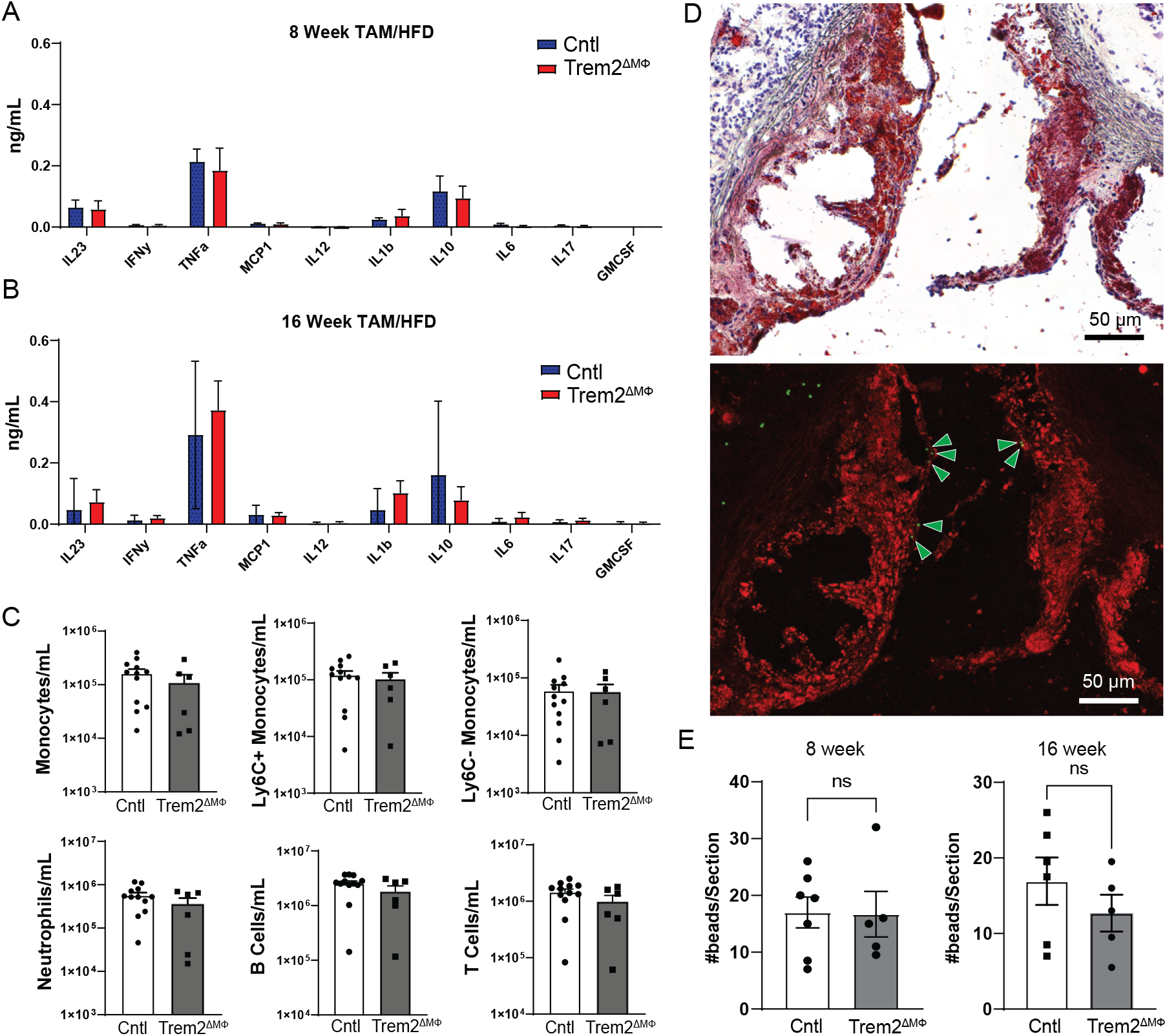
Conditional deletion of Trem2 has no effect on monocyte recruitment or systemic inflammation. A) Serum from 8 week TAM-HFD fed mice were assessed for cytokine levels by multiplex assay. B) Serum from 16 week TAM-HFD fed mice were assessed for cytokine levels by multiplex assay. C) Blood immune cells were assessed after 16 weeks TAM-HFD by flow cytometry. D) Monocyte recruitment was assessed by bead labeling and recruitment experiment, images from representative histologic and immunofluorescence image with lipid content (red) and beads (green). E) Quantification of plaque-associated beads that were counted per section for 8 or 16 week TAM-HFD experiments.

We next performed a monocyte recruitment assay by labeling monocytes with fluorescent beads to determine whether there were changes in recruitment to lesions following 8 or 16 weeks of TAM-HFD. Following established protocols^53,54^, beads were injected i.v. to label classical monocytes, labeling efficiency was checked after 24 hours, and mice were sacrificed for monocyte infiltration into lesions 48 hours after bead labeling. Figure 5D shows representative plaque area and bead recruitment into the lesion. Beads typically infiltrate the surface of lesions, as previously described^54^. Importantly bead uptake by blood monocytes was similar between control and Trem2^ΔMФ^ mice (Supplemental Figure 4C). Quantification of beads in atherosclerotic lesions revealed comparable recruitment rates between experimental groups (Figure 5E). This was independent of changes in atherosclerotic lesion size, suggesting that changes in plaque size were likely associated with local changes in foamy macrophage function or persistence in tissue.

To test local plaque changes in TAM-HFD fed Trem2^ΔMФ^ or control mice, we performed confocal microscopy to assess plaque macrophage and smooth muscle area using immunofluorescence staining of aortic root sections. Macrophages were identified using CD68 antibody and smooth muscle cells with alpha actin (SMA) (Supplemental Figure 5A-B). Quantification confirmed reduced macrophage area in 8 week and 16 week treatment groups (Supplemental Figure 5 C-D). However, percentage of plaque area was skewed toward greater macrophage representation in Trem2^ΔMФ^ plaque compared to controls. SMA coverage was not detectably different between groups at either time point. The enhanced macrophage representation relative to area is consistent with a less mature atherosclerotic plaque, whereas the larger control plaques may begin developing necrotic cores due to advanced disease state.

Next, to detect potential changes in proliferation, we performed immunostaining for Ki67 (Figure 6A). Quantification of CD68^+^ Ki67^+^ macrophages within lesions showed a dramatic reduction in local proliferation in Trem2^ΔMФ^ plaques at both 8 and 16 week TAM-HFD feeding (Figure 6B). To test whether loss of Trem2 resulted in changes in macrophage susceptibility to death in plaques, we also performed TUNEL staining to identify apoptotic cells (Figure 6C). TUNEL^+^ CD68^+^ macrophages were dramatically enriched in Trem2^ΔMФ^ plaques at both time points analyzed (Figure 6D). Together, these data suggest that foamy macrophages rely on Trem2 to persist and proliferate in atherosclerotic lesions.

**Figure 6.**
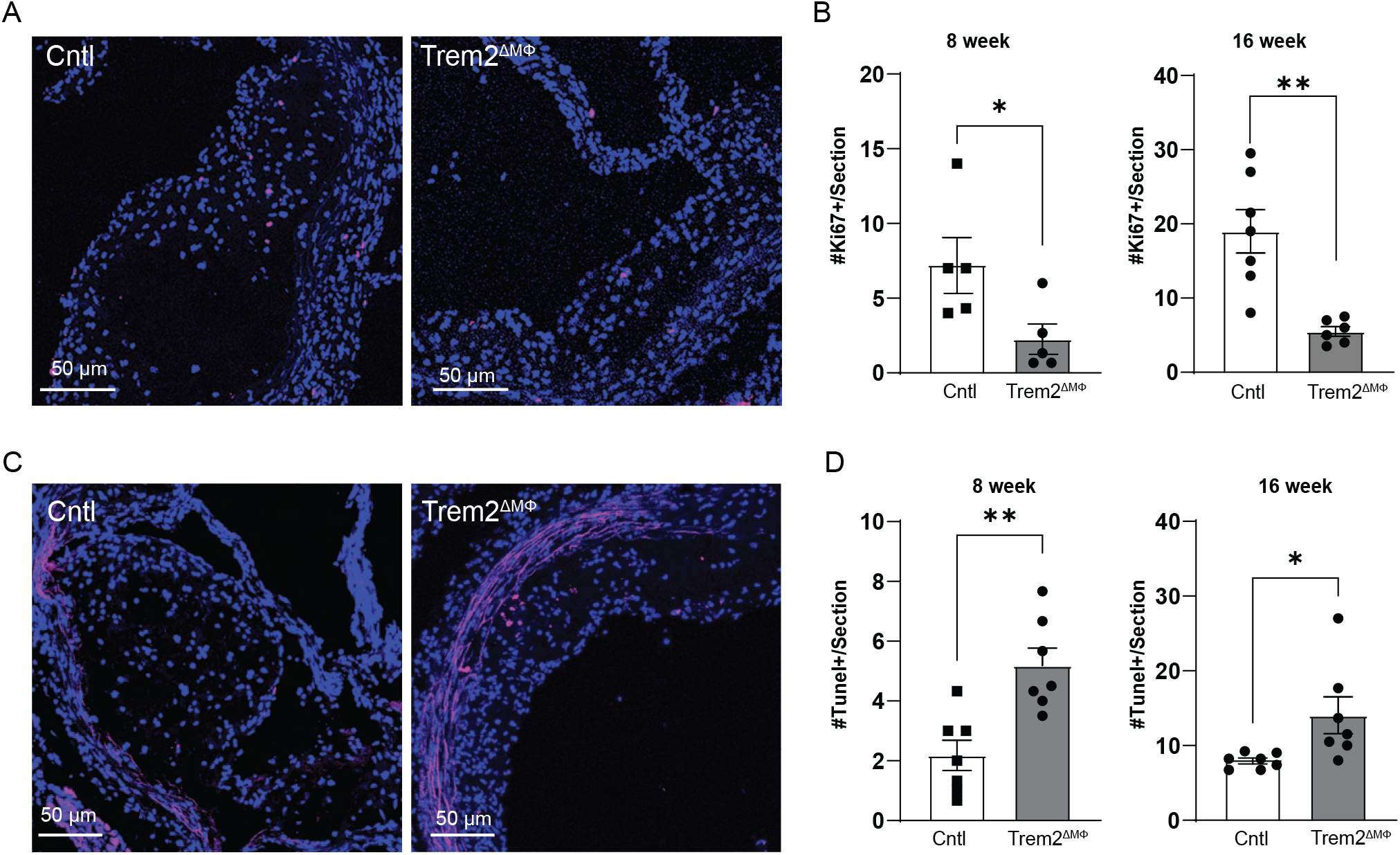
Trem2 regulates foamy macrophage survival and proliferation in atherosclerotic lesions. A) Confocal micrograph showing Ki67 staining for proliferation in control (Cntl) or Trem2-deficient mice after 16-weeks TAM-HFD feeding. B) Quantification of Ki67+ macrophages (CD68+) per section in 8- or 16-week TAM-HFD samples. C) Confocal micrograph of TUNEL staining for detection of dying cells within atherosclerotic lesions after 16-weeks TAM-HFD feeding. D) Quantification of TUNEL+ macrophages (CD68+) per section in 8- or 16-week TAM-HFD samples.

### Trem2 deletion in establish plaque slows atherosclerosis progression

Since humans are diagnosed with atherosclerosis once it has already developed, we wanted to test whether Trem2 can be targeted therapeutically in established atherosclerotic lesions. Therefore, we designed an in vivo intervention study by feeding Trem2^ΔMФ^ mice or littermate controls a regular HFD for 8 weeks to induce atherosclerotic lesions in all animals, then transitioning the mice to a TAM-HFD diet to allow for deletion of Trem2 on CX3CR1-expressing cells for an additional 8 weeks of time. After 16 weeks of total HFD feeding, mice were sacrificed and assessed for atherosclerosis progression (schematic in Figure 7A). Aortic arch and aortic sinus were measured for atherosclerotic plaque area and we found that deletion of Trem2 on myeloid cells in established lesions attenuated further atherosclerosis progression (Figure 7B-C). This outcome was independent of blood monocyte numbers or serum cholesterol levels (Figure 7D-E). Similar to data from the continuous treatment experiments, plaque macrophages from Trem2^ΔMФ^ mice showed elevated TUNEL and reduced Ki67 expression compared to controls (Figure 7F-G). Overall, these data emphasize the potential for targeting Trem2 to reduce further atherosclerosis progression in patients with established vascular plaques.

**Figure 7.**
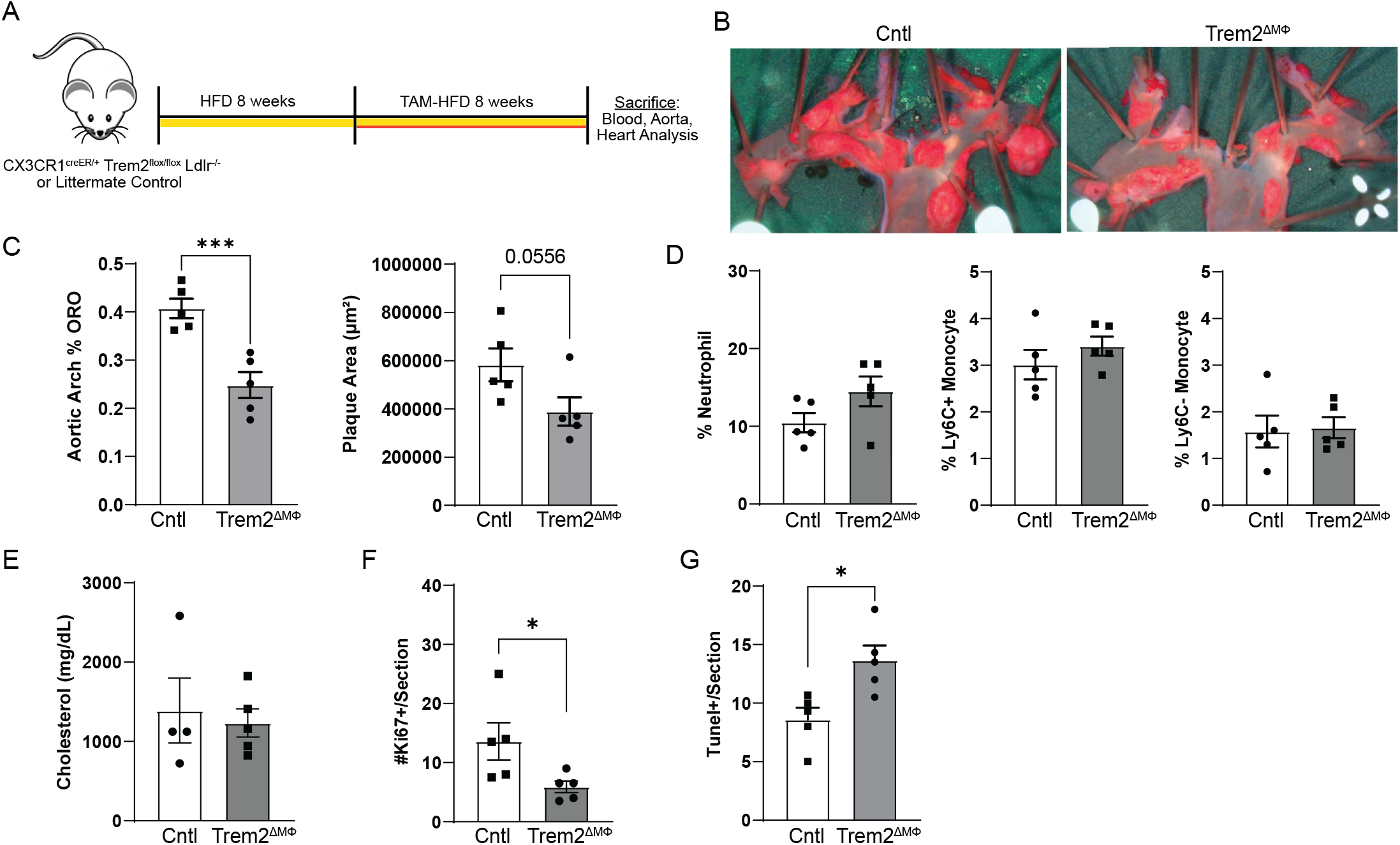
Deletion of Trem2 in established atherosclerotic lesions leads to enhanced foamy macrophage death and reduced atherosclerotic plaque size. A) Schematic for intervention study where mice were fed HFD for 8 weeks, then switched to TAM/HFD for an additional 8 weeks before sacrifice. B) En face aorta analysis of plaque area after 16 weeks diet-switch intervention study. C) Quantification of plaque area in aorta and aortic sinus. D) Blood immune population analysis after 16-week diet switch intervention model. E) Total serum cholesterol levels after 16-week diet switch model F) Quantification of plaque macrophage proliferation analysis by Ki67+ macrophages (CD68+). G) Quantification of TUNEL+ macrophages (CD68+) in plaques after 16-week diet switch model.

### Trem2 regulates foamy macrophage cholesterol sensing and ER stress response

To further examine the mechanisms that are regulated downstream of Trem2 in foamy and nonfoamy macrophages, we generated a Trem2^-/-^ BV2 macrophage cell line. For in vitro studies, BV2 cells were cultured in media alone, in the presence of soluble cholesterol at 20 μg/mL, or “high” dosing at 80 μg/mL. First, we validated that we had CRISPR-Cas9 knockout of Trem2 protein in BV2 cells following soluble cholesterol treatment (Figure 8A). We next sought to understand the molecular regulation of Trem2 in nonfoamy and foamy macrophages. Thus, we performed bulk RNA-seq analysis of control or Trem2^-/-^ BV2 cells treated overnight in media alone or 20 μg/mL soluble cholesterol. We were interested in understanding the response to cholesterol loading, so we compared WT BV2 to WT-foamy and Trem2^-/-^ BV2 to Trem2^-/-^-foamy. As expected, control (WT) foamy treated cells showed a strong upregulation of cholesterol efflux genes (*Abca1, Abcg1*), and a downregulation of cholesterol synthesis genes (*Cyp51, Hmgcr*) (Figure 8B). Surprisingly, these features were inversely associated in the Trem2^-/-^ foamy cells. Cholesterol loaded Trem2^-/-^ cells did show a lipid loading phenotype by increased expression of *Fabp5, Stard4*, and *Plin2*, but actually had reduced expression of efflux (*Abca1, Abcg1*) and upregulation of cholesterol synthesis genes (*Cyp51, Hmgcr*) (Figure 8C). Comparison between WT and Trem2^-/-^ macrophages (untreated or foamy) revealed numerous classical and alternative activation pathways being upregulated in Trem2^-/-^ BV2, independent of treatment condition (Supplemental Figure 6A-B). In addition, cell cycle genes, including *Ccnd1* and *Ccnd2*, were upregulated in WT BV2 cells.

**Figure 8.**
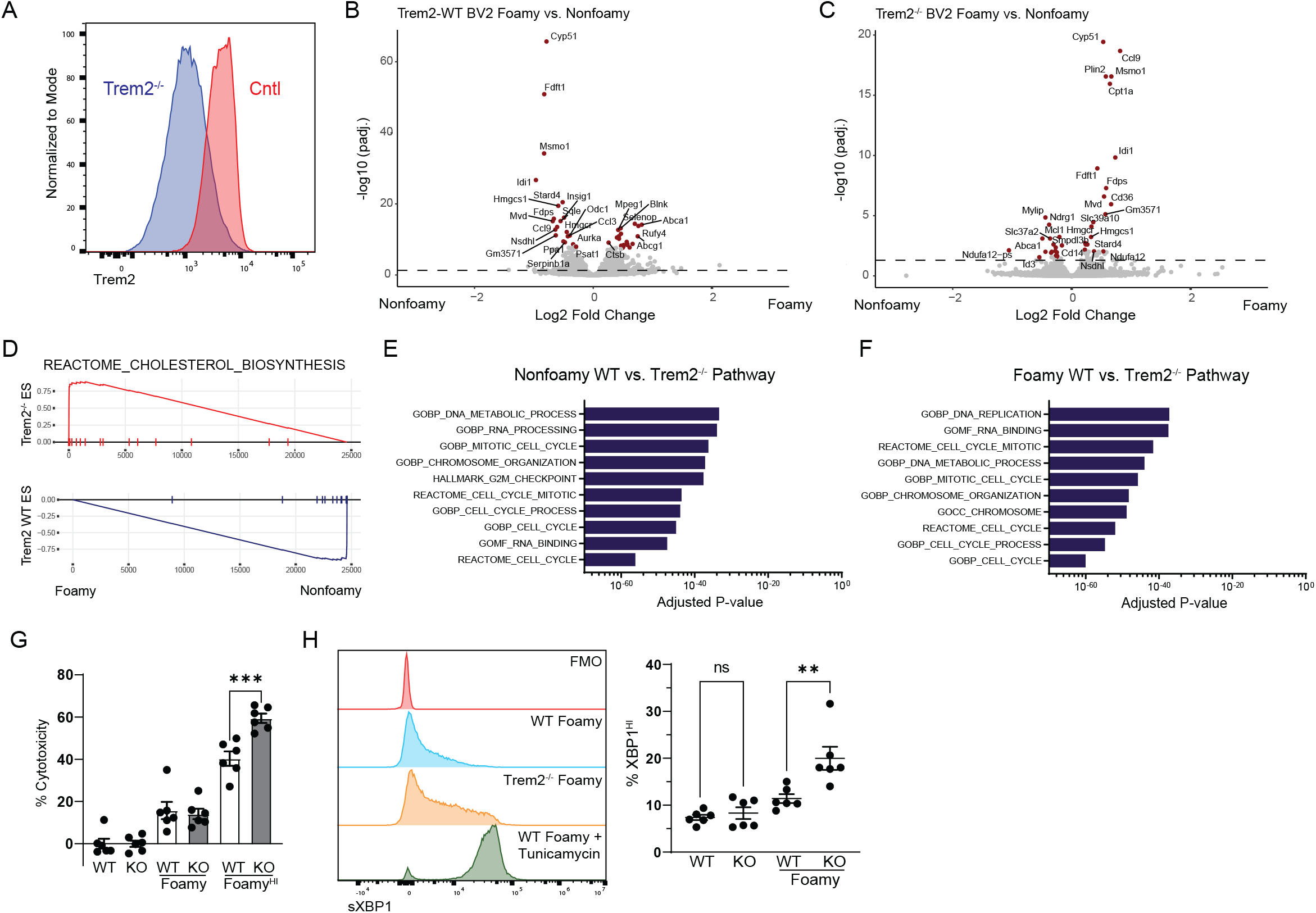
Trem2 deficient foamy macrophages are susceptible to cell death and enhanced ER-stress response. A) Trem2^-/-^ BV2 macrophages were developed using CRISPR/Cas9 deletion approach and differentiated into foamy macrophages by soluble cholesterol treatment. Cells were treated overnight and then assessed for Trem2 expression by flow cytometry. B) WT or (C) Trem2^-/-^ BV2 macrophages were differentiated in media control or media with 20 μg/mL soluble cholesterol overnight, then bulk sequenced by RNA-seq. D) GSEA plot of Cholesterol Biosynthesis pathways. E) Pathway analysis of RNA-seq data comparing WT and Trem2^-/-^ nonfoamy BV2 cells. E) Pathway analysis of RNA-seq data comparing WT and Trem2^-/-^ foamy BV2 cells. G) WT or Trem2^-/-^ BV2 macrophages were differentiated in media control, media with 20 μg/mL or 80 μg/mL soluble cholesterol to induce foamy macrophage formation. Cell supernatant was assessed for cytotoxicity by LDH assay after 16 hours. H) WT or Trem2^-/-^ BV2 macrophages were differentiated in media control or media with 20 μg/mL soluble cholesterol, then assessed for activation of ER stress response by sXBP1 levels by flow cytometry. Tunicamycin was used as a positive control. (right) Quantification of ER stress response in control or foamy induced WT and Trem2^-/-^ BV2 macrophages.

Consistent with our interpretation of dysfunctional cholesterol sensing and response, Trem2^-/-^ BV2 showed upregulation of cholesterol biosynthesis pathways, whereas WT cells showed significant downregulation, as observed by GSEA plot (Figure 8D). Pathway analysis comparing WT-foamy and Trem2^-/-^-foamy macrophages revealed that the top 10 pathways enriched in WT nonfoamy and foamy macrophages were directly associated with cell cycle pathways (Figure 8E-F, Supplemental Figure 6C), supporting our prior conclusion that Trem2 was required for foamy macrophage proliferation. Overall, RNA-seq analysis revealed dramatic changes in cell proliferation and lipid metabolism pathways in Trem2^-/-^ BV2 cells compared to WT.

Since gene expression analysis suggested a defect in Trem2^-/-^ BV2 cells in cell cycle and lipid sensing, we first tested whether these cells were more sensitive to lipid loading. Using an LDH assay, we tested cytotoxicity in foamy and nonfoamy BV2 KO and WT clones loaded with soluble cholesterol (Figure 8G). Using soluble cholesterol bypasses any defects associated with Trem2^-/-^ lipid loading and forces cell accumulation of cholesterol. Data show no change in Trem2^-/-^ BV2 cell cytotoxicity until cells were pushed with greater loads of soluble cholesterol (Foamy^Hi^), whereas Trem2^-/-^ foamy cells showed approximately a 20% increase in overall death in culture, supporting observations from in vivo plaque analysis.

Since Trem2^-/-^ foamy macrophages showed enhanced cytotoxicity and defective cholesterol response profile, we hypothesized that loss of Trem2 in foamy macrophages may lead to accumulation of free cholesterol in the endoplasmic reticulum (ER) and promote the ER stress response. This is supported by prior work showing that Trem2-deficient microglia were unable to adapt to excess cholesterol exposure and form additional lipid droplets following myelin debris uptake^55^. Cholesterol mediated cytotoxicity is commonly associated with an ER-stress responses, so we performed overnight foamy macrophage formation assay in BV2 cells.

We then performed intracellular flow cytometry for intracellular sXBP1, the active form of the protein that drives transcription of ER stress response genes (Figure 8H)^56^. Tunicamycin treatment was used as a positive control, which showed comparable levels between WT and Trem2^-/-^ BV2 cells regardless of foamy status. In our experimental group, we observed enhanced ER stress response in Trem2^-/-^ foamy macrophages after 20 μg/mL soluble cholesterol, even when there was no difference in cell death (Figure 8H). Together these data support a model where Trem2 regulates intracellular lipid sensing and metabolic programming in foamy macrophages which allows for promotion of foamy cell survival and maximization of lipid storage and efflux potential.

## Discussion

Macrophages are major contributors to the formation of atherosclerotic plaque. Many features of lipid loaded foamy macrophage function are well-established, including cholesterol uptake, storage, and efflux. However, factors specifically regulating foamy macrophage differentiation and survival have remained understudied. We approached this subject using an in silico trajectory analysis approach of scRNA-seq data. By generating a META-plaque scRNA-seq dataset, we were able to achieve (i) finer resolution to identify rare clusters of cells (such as monocytes), (ii) observe previously concealed intermediate cells between clusters, and (iii) split previously-defined clusters into sub-clusters based on gene expression. META scRNA-seq datasets have been generated in prior studies, but the majority of these studies have focused on describing cellular diversity through the formation of an atlas defining cluster identities^9,45^. In addition, the use of computational tools to predict differentiation trajectories between cell clusters with known biological relationships has been lacking in the cardiovascular field. In our prior work, we hypothesized the trajectory of monocytes entering atherosclerotic lesions, suggesting that monocytes may make commitment decisions directly upon entry into lesion microenvironments^57^. For our current study, we tested this possibility to understand the differentiation trajectory between major cluster subsets. By gaining a clearer picture of cluster populations and intermediaries, we were able to define gene expression trajectories associated with the major inflammatory or foamy macrophage states that we predict are terminal differentiation points. In the case of foamy cell formation, it is interesting that genes appear to turn on in stages as they progress along pseudotime. Early commitment genes include *Trem2, Fabp5, Plin2*, and *Lgals3*, whereas *ATP6v0d2* is not induced until late stages of commitment.

One potentially significant result from our analysis was the predicted binary differentiation pathways from recruited monocytes toward inflammatory and foamy macrophage lineages. Studies in other chronic inflammatory disease models, including lung fibrosis, suggest that monocytes undergo a transient inflammatory state before slowly maturing into a proresolving tissue resident-like macrophage^58^. Follow up studies are needed to test this hypothesis by utilizing modern fate-mapping and spatial mapping approaches to determine whether foamy and inflammatory clusters are indeed terminal differentiation states, or whether there is plasticity between the major plaque associated clusters. Given the highly inflammatory state of Cluster 1 defined in Figure 1C, it may not be surprising if many of these cells fail to persist for long periods of time due to inflammasome activation and subsequent pyroptosis^59,60^. If true, the MHCII+ Cluster that lacks inflammatory signature may be a rare cell that survives inflammasome activation within lesions, then progresses into a nonfoamy plaque resident macrophage. Expanded investigation into these possibilities and defining the function of the newly described MHCII+ subsets cells will be of great interest to the field moving forward.

Pseudotime analysis was used to identify genes associated with commitment toward foamy or inflammatory outcomes. This revealed an importance index of predicted genes that were strongly associated with differentiation outcomes. We also performed an unbiased genome-wide CRISPR screen to detect genes that regulate the uptake of oxLDL. This approach was performed in the presence of pre-formed foamy macrophages, thus replicating features of cells that are previously differentiated into foamy macrophages. Together, the screen and pseudotime analysis emphasized the potential importance of Trem2 in foamy cell formation. Through in vitro and in vivo approaches, we validated the role of Trem2 in regulating foamy macrophage lipid, cellular metabolism, and survival in lesions. Furthermore, gene expression analysis identified pathways that were dysfunctional in Trem2-deficient foamy macrophages, including cholesterol synthesis and efflux pathways.

Whole body Trem2^-/-^ mice possess a wide variety of phenotypes that might make analysis and interpretation of the role of Trem2 difficult, particularly in the context of atherosclerosis or other chronic diseases. Relevant to our study is the observation that Trem2^-/-^ mice have exacerbated development of metabolic disease. Multiple studies have found that Trem2 signaling on adipose macrophages promotes lipid-associated macrophage homeostasis, and whole body Trem2^-/-^ mice have elevated serum cholesterol, increased systemic proinflammatory cytokines, and increased body weight accumulation upon HFD feeding, driven by massive adipocyte hypertrophy^27,61^. Moreover, mice lacking Trem2 have also been found to develop accelerated fatty liver disease and steatohepatitis due to increased inflammation, further emphasizing systemic alterations in this model^28,62^. Interestingly, we report that temporal deletion of Trem2 specifically on macrophages, using the CX3CR1^creER^ Trem2^flox^ Ldlr^-/-^ model, does not recapitulate the systemic effects seen with whole body Trem2^-/-^. This leads us to hypothesize that differences between models could be due either to developmental defects that require Trem2, such as in brain or liver, or another possibility is that Trem2 deletion influences cell function in non-macrophages subsets, as Trem2 expression is not restricted to only macrophages^63,64^. Despite differences seen between these models, our findings align with the aforementioned studies and support the idea that Trem2 is a master regulator of lipid associated macrophage (LAM) function and phenotype across disease subtypes.

Mechanistically, we found that Trem2 signaling promotes proliferation and survival of foamy macrophages. Deletion of Trem2 in foamy macrophages led to a defect in upregulation of cholesterol efflux and downregulation of cholesterol synthesis pathways, which supports a lipid sensor role for Trem2. This result is consistent with other groups who have also reported deficiencies in cholesterol efflux pathways in Trem2-deficient macrophages^14,65^. Our studies further revealed an increase in ER stress responses in Trem2^-/-^ foamy macrophages, potentially driving cell death upon lipid loading. ER stress responses may have been largely overlooked in models that drive less severe lipid content in macrophages, such as Alzheimer’s disease. In line with our results, previous studies in microglia have found that Trem2 deletion leads to impaired uptake and storage of myelin debris into lipid droplets^55^. We hypothesize that similar mechanisms are at play in the setting of excess cholesterol, where Trem2-deficient macrophages cannot effectively store cholesterol into droplets, and instead accumulate lipids to the ER, leading to excess stress responses and ultimately cell death^55^. It is possible that excess ER stress responses are linked to metabolic reprogramming through generation of reactive oxygen species (ROS) that accumulate during cellular stress. Transcription factors such as HIF1a has been extensively shown to be driven by macrophage ROS production^66^ and can directly promote a metabolic switch favoring glycolysis^67^. Based on this, we hypothesize that the excess ER stress response found following Trem2 deletion could lead to an inability of Trem2-deficient foamy macrophages to switch toward oxidative phosphorylation metabolism and are likely dependent on glycolysis, but further mechanistic studies are needed to elucidate this possible interplay.

Trem2 has been proposed for a therapeutic target for a variety of disease models, including Alzheimer’s disease^68^ and cancer^69^. Targeting Trem2 is temping because of its immunomodulatory function, yet it is not associated with a major role in acute inflammation such as infection. Prior approaches to target inflammation in cardiovascular disease, such as IL1β blockade, have failed during clinical trials, primarily due to broad immunosuppressive effects leading to impaired host defense against microbial infection^70^. Given this, we suggest that targeting Trem2 in atherosclerosis would allow for effective modulation of plaque macrophages without impairing responses to infection. Importantly, both Trem2 agonistic and blocking antibodies have been extensively tested in both mice and humans for use in Alzheimer’s disease and cancer. Trem2 blocking antibodies have been shown to enhance tumor immunotherapy action through modulating the cancer microenvironment^69^. Similar approaches could be used in atherosclerosis which may allow for more efficient approaches to reduce inflammatory signatures in disease. However, in Alzheimer’s disease, Trem2 agonism using monoclonal antibodies has been shown to promote proliferation and survival of microglia while decreasing pathogenic local proinflammatory responses, leading to improved outcomes^22^. Despite our findings that Trem2 deletion in mice leads to decreased plaque sizes, we propose that targeting Trem2 agonistically in atherosclerosis could be potentially advantageous by promoting macrophage survival and local anti-inflammatory responses which would improve unfavorable plaque phenotypes such as necrotic core formation that is often found in advanced human disease. Moreover, given the proposed role for Trem2 in mediating cholesterol efflux downstream of LXR activation^14,71^, utilization of Trem2 agonists in atherosclerosis could drive efflux of cholesterol from lipid laden macrophages and reinvigorate pro-efferocytotic functions to clear dying cells. Thus, we suggest that translational efforts to block Trem2 in atherosclerosis should also consider accompanying approaches to promote Trem2 signaling in advanced diseased settings rather than simply impair its function.

In conclusion, we have identified Trem2 to be enriched on foamy macrophages, but not inflammatory macrophages, in atherosclerotic plaques. Loss of Trem2 in foamy macrophages led to enhanced cellular stress response, reduced proliferative potential, and augmented cell death. Conditional deletion experiments showed that Trem2 was required in foamy macrophages for atherosclerosis progression, and that even targeting Trem2 in established lesions was sufficient to reduce overall plaque burden. Thus, Trem2 is a novel regulator of foamy macrophage formation and function, and is an appealing target for future therapeutic intervention studies.

## Acknowledgements

We would like to thank Dr. Bruce Lamb (Indiana University) for kindly sharing the Trem2^flox^ mouse and Dr. Marco Colonna (Washington University in St. Louis) for kindly sharing the Trem2^-/-^ mouse strain. We would like to thank Dr. Maxim Artyomov (Washington University in St. Louis) for access to his scRNA-seq portal. We would like to thank the UMN anatomy bequest program and families of donors for artery samples to advance education and science. We would also like to thank the University of Minnesota Flow Cytometry core facility and UMN Center for Immunology Imaging core for assistance in data acquisition. Research was funded by grant support from R01 AI165553 (JWW), R00 HL138163 (JWW), American Heart Association (AHA) CDA855022 (JWW), and Minnesota Office of Higher Education Award (JWW). MTP was supported by a predoctoral research fellowship AHA 903380.

## Author Contributions

JWW, MTP, XR, IS, TB and KZ conceived and designed the research project. MTP, MF, YX, CB, AZ, PRS, CER, GF, AEK, NA, KZ, JWW performed experiments. MTP and JWW wrote the manuscript and prepared figures. All authors assisted with data interpretation and manuscript revisions.

## Supplemental Figure Legends

**Supplemental Figure 1. Integrated scRNA-seq differential gene expression**

Heat map of cell clusters identified in META-scRNA-seq dataset, and top enriched genes for each cluster.

**Supplemental Figure 2. Trajectory analysis of META-scRNA-seq dataset**

A) Trajectory analysis revealed stages of differentiation following monocytes entry associated with inflammatory macrophage clusters

B) Upregulation of inflammation genes associated with entry into intima and commitment toward inflammatory macrophage differentiation

C) Ccl3 and Ccl4 were uniquely expressed by committed inflammatory cells, but not by intermediate inflammatory cells.

D) MHC II expression is gradually elevated along commitment toward inflammatory differentiation arm of the trajectory map.

**Supplemental Figure 3. CRISPR screen of foamy macrophage oxLDL uptake**

A) CRISPR guide enrichment by rank-order was plotted against P-value for DiI-oxLDL-low compared against DiI-oxLDL-high to identify top enriched guides.

B) Top 15 “importance index” genes associated with foamy cell commitment by Trade-seq analysis (Figure 1), were compared for gene rank and enrichment in CRISPR screen.

**Supplemental Figure 4.**

A) Flow cytometric gating strategy for identifying major blood immune cell populations.

B) Blood immune cell profiling by flow cytometry in indicated mice after 16 weeks TAM-HFD feeding.

C) Classical monocyte bead uptake in the blood was measured by flow cytometry 24 hours after i.v. bead injection in indicated strains after 16 weeks TAM-HFD feeding.

**Supplemental Figure 5. Macrophage and smooth muscle quantification in plaque**

A) Representative imaging of macrophages in plaque, by CD68 staining

B) Representative imaging of smooth muscle cells, by SMA staining.

C) 8-week and (D) 16-week analysis of plaque macrophage area and percentage coverage with smooth muscle (SMA) or macrophage (CD68).

**Supplemental Figure 6. Bulk RNA-seq analysis of WT or Trem2^-/-^ BV2 cells**

A) Heat map of nonfoamy macrophages comparing top enriched WT and Trem2^-/-^ genes.

B) Heat map of foamy macrophages comparing top enriched WT and Trem2^-/-^ genes.

C) Normalized enrichment scores for top pathways associated with WT BV2 compared to Trem2^-/-^ BV2 in media alone (nonfoamy) or following foamy differentiation.

