## Supplemental Figures for "Trem2 Promotes Foamy Macrophage Lipid Uptake and Survival in Atherosclerosis"

### SUPPLEMENTAL FIGURE 1

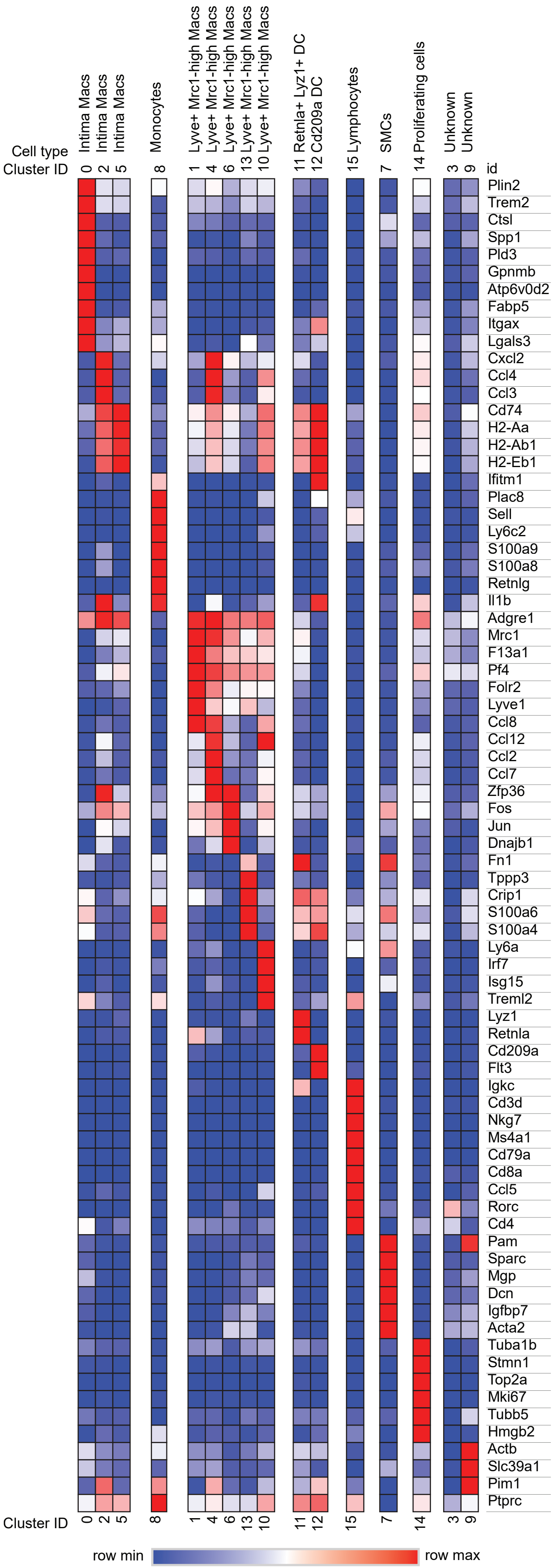

### SUPPLEMENTAL FIGURE 2

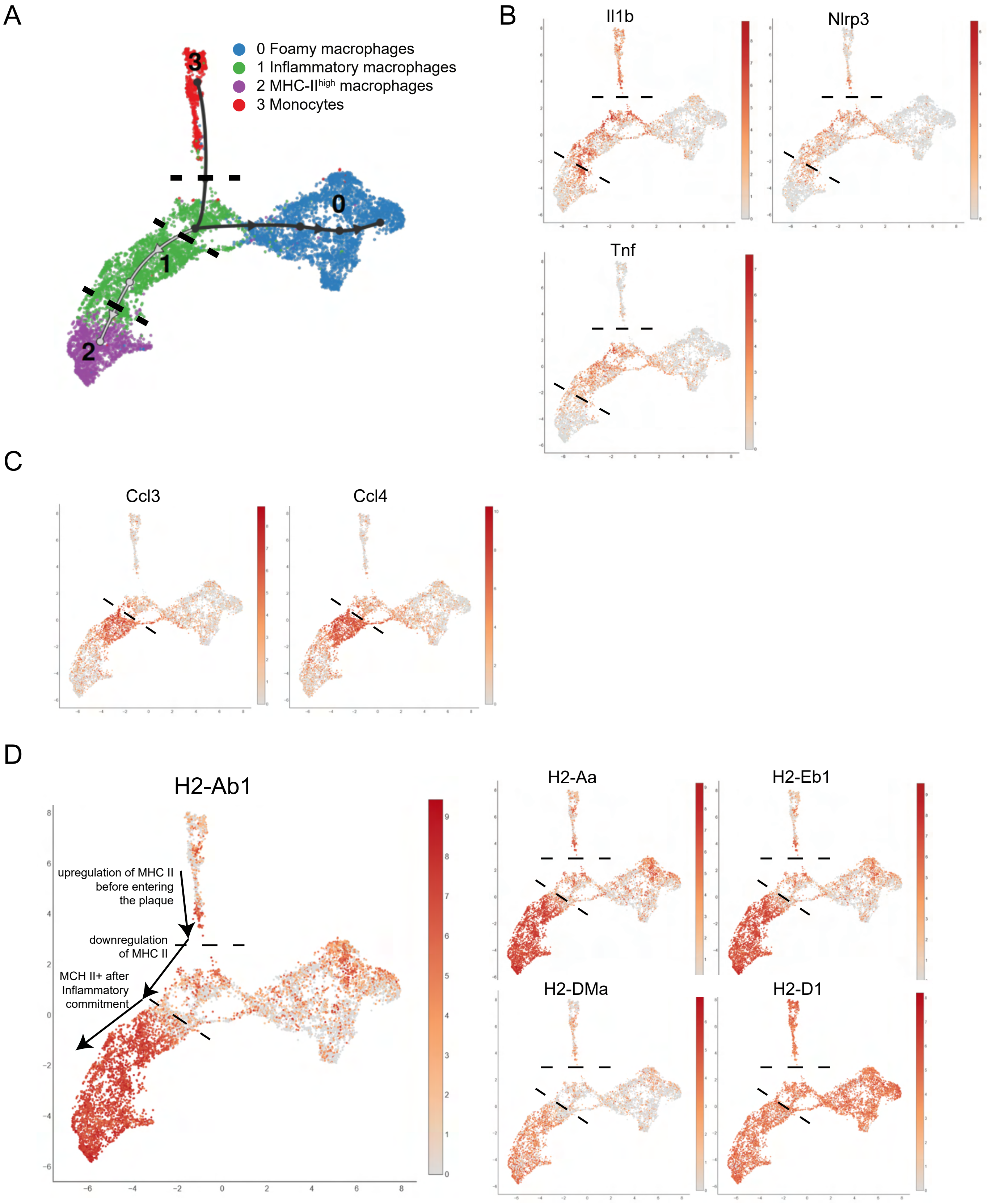

SUPPLEMENTAL FIGURE 3

A

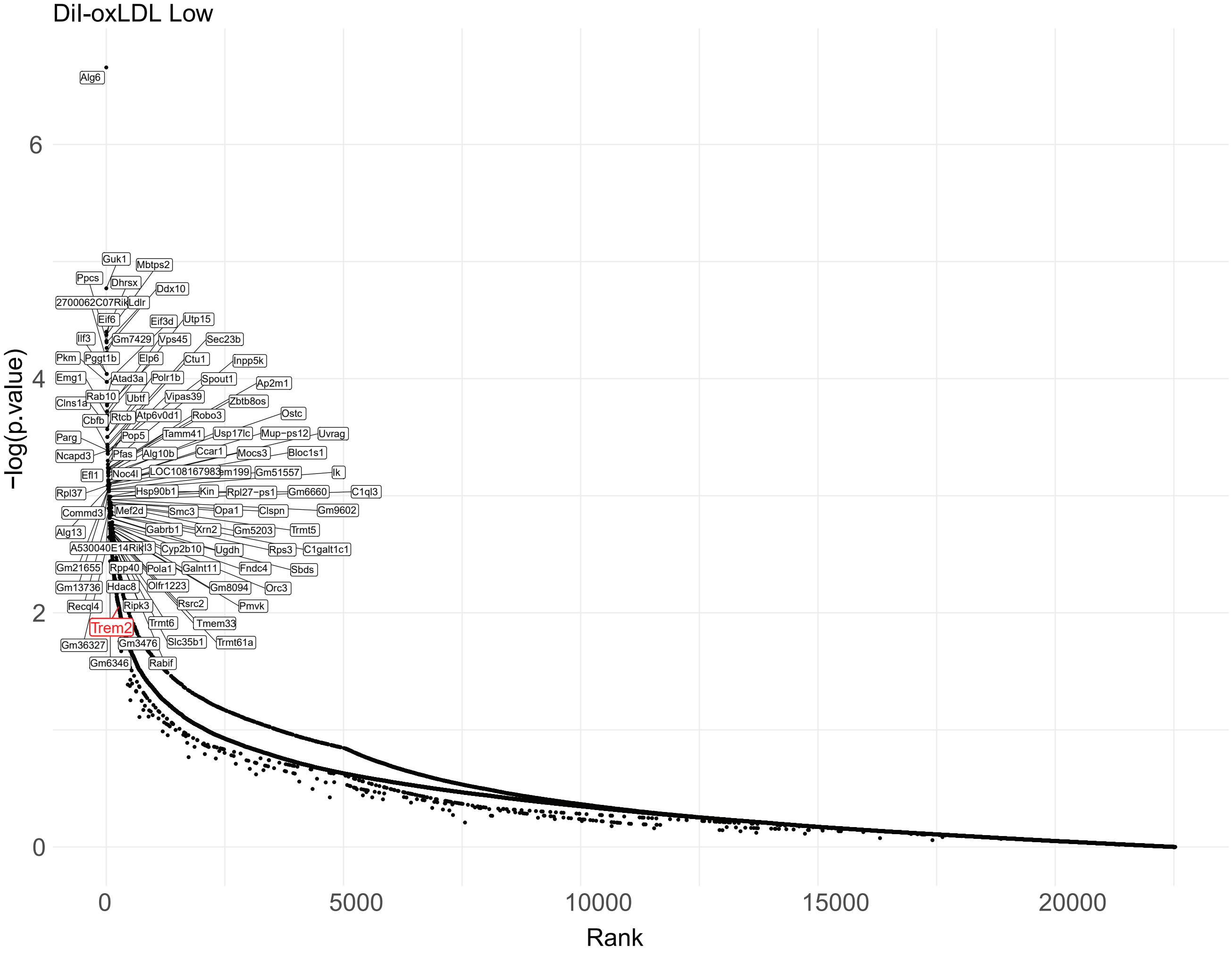


B

| ID | Gene Rank | Pos. Score | P-value | Log FC |
|---|---|---|---|---|
| Lgals3 | 13140 | 0.587 | 0.588 | 0.320 |
| Spp1 | 7269 | 0.280 | 0.332 | 0.192 |
| Trem2 | 267 | 0.005 | 0.009 | 1.820 |
| Cd9 | 9843 | 0.423 | 0.442 | 0.670 |
| Ctsb | 6784 | 0.251 | 0.311 | -0.054 |
| Cxcl10 | 3274 | 0.089 | 0.154 | 0.515 |
| Kdm6b | 9049 | 0.381 | 0.408 | -0.334 |
| Cxcl12 | 12966 | 0.579 | 0.580 | -0.086 |
| Lpl | 6136 | 0.213 | 0.284 | 0.197 |
| Ctsd | 12328 | 0.550 | 0.552 | -0.118 |
| Cd63 | 15068 | 0.677 | 0.677 | 0.140 |
| Ccl4 | 3677 | 0.102 | 0.173 | 0.835 |
| Capg | 20053 | 0.892 | 0.892 | -0.747 |
| Cxcl1 | 5516 | 0.174 | 0.257 | 0.714 |
| Marcksl1 | 16753 | 0.749 | 0.749 | -0.698 |

SUPPLEMENTAL FIGURE 4

A

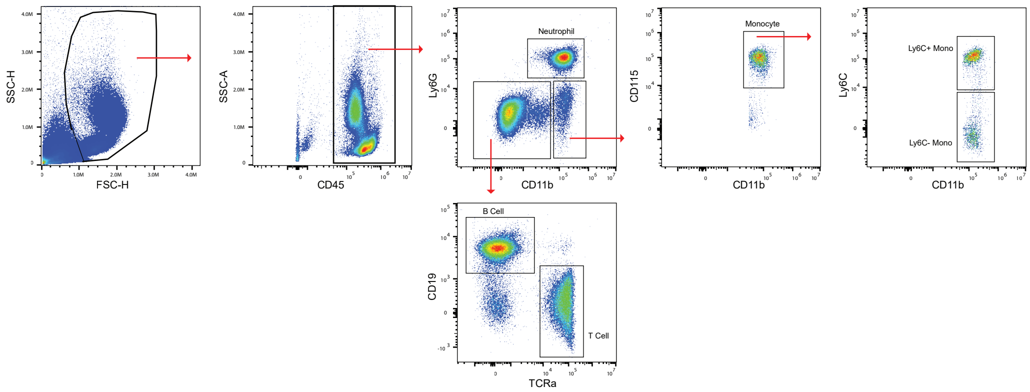


B

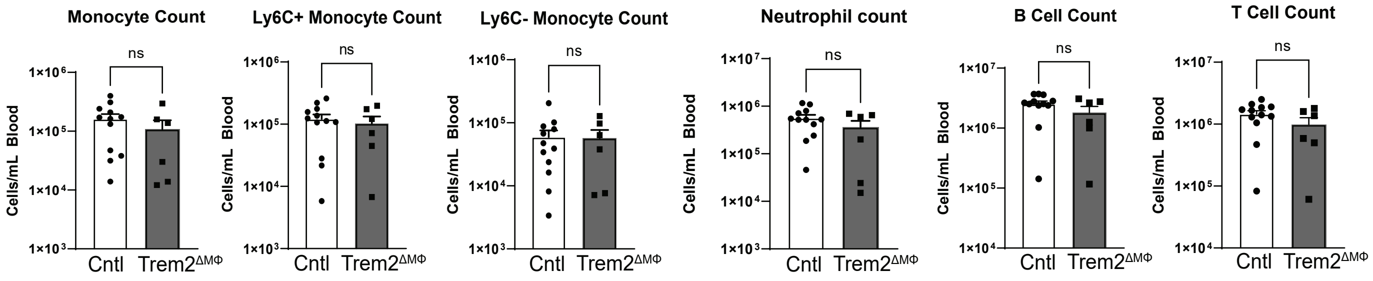


C

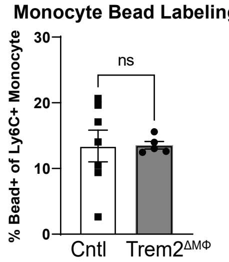

SUPPLEMENTAL FIGURE 5

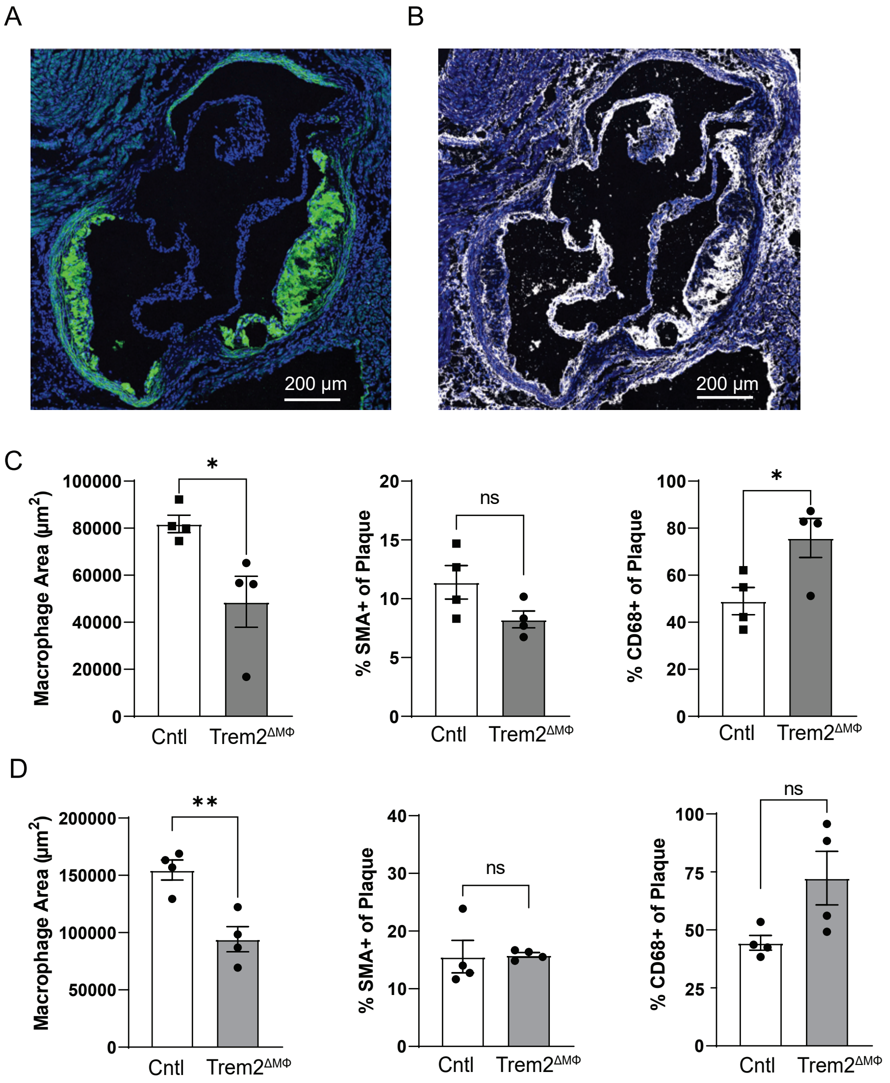

SUPPLEMENTAL FIGURE 6

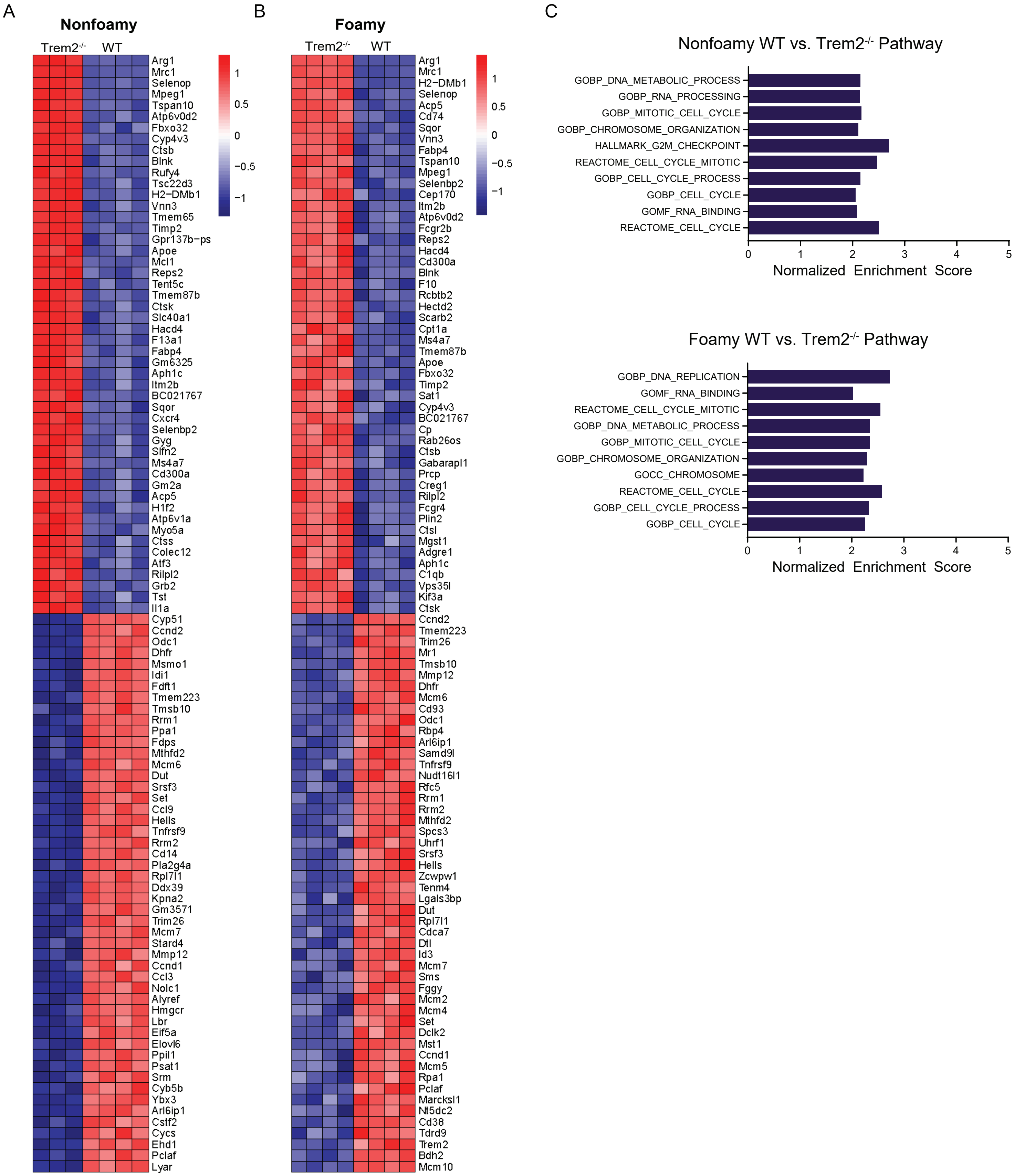
